# Probing Genomic Foundation Models with Splice-Variant Perturbations

**DOI:** 10.64898/2026.10.02.756289

**Authors:** Marcel V. Alavi

## Abstract

As benchmarks saturate, traditional validation methods fail to capture the probabilistic representations of genomic foundation models. This study probes self-supervised generalists (NTv3 650M, Evo2 40B, Genos 10B) alongside task-specific specialists (SpliceAI, AlphaGenome, Borzoi) using a mechanistically focused paradigm centered on expert-curated single-nucleotide substitutions within the splicing regions of the exon-rich *OPA1* gene. Extending this analysis across the broader corpus of *OPA1* mutations, variants of uncertain significance (VUS), and an independent 65-gene dataset of spliceogenic mutations demonstrates that foundation models not only match task-specific networks, but also generalize across mutation types. Training data scale, genetic diversity, and evolutionary context are key for capturing RNA splicing syntax, whereas sparse routing may constrain performance. Mechanistically, foundation models exhibit a compute asymmetry: Evo2 40B frontloads compute into massive parameter scale to resolve single-nucleotide variants within a narrow 425-bp context window, whereas the bidirectional NTv3 650M model and the Genos 10B mixture-of-experts (MoE) architecture backload compute to inference, requiring spatial logit aggregation and vast context windows of up to 94 kb. Furthermore, accuracy may benefit from locus-specific thresholding rather than mutation-class calibration, exposing representation boundaries of the self-supervised models.

## I. INTRODUCTION

**A**ccording to the National Human Genome Research Institute, we share about 99.6% sequence identity with the human reference genome. The remaining 0.4% variation determines individual phenotypes and disease susceptibility beyond altering protein-coding sequences. Importantly, genomic variants regulate RNA expression, transcript stability, and alternative splicing.

Characterizing this complex regulatory syntax remains challenging, but computational tools now play a vital role in interpreting genetic variant effects. For instance, sequence-to-function models ingest raw DNA sequences to predict functional outputs without relying on prior annotations or sequence homology. Instead, they map individual base pairs to experimental signals like read counts, epigenetic marks, or chromatin accessibility. Historically, these discriminative models were optimized for single classification or regression tasks, computing the conditional probability *P*(*Y*|*X*) to draw a decision boundary between phenotype classes *Y* given genotype *X*. Today, high-throughput datasets allow the training of genomic foundation models via self-supervised learning. Instead of just drawing boundaries, these architectures model the underlying sequence distribution *P*(*X*) or joint probability *P*(*Y, X*). Consequently, they move beyond static predictions to simulate diverse biological outcomes or generate completely novel sequences.

However, traditional validation methods fail to capture the nuanced, probabilistic outputs of these models. Standard benchmarks have largely saturated and are often compromised by data leakage during pretraining, which makes them unreliable indicators of real-world utility. To address this gap, the present study appraised a mechanistically focused testing paradigm centered on single-nucleotide substitutions within the splicing regions of an individual, exon-rich gene.

Autosomal Dominant Optic Atrophy (ADOA) is a rare form of blindness primarily caused by *OPA1* mutations identified in 10% to 37% of patients with optic atrophy [1-4]. The human *OPA1* gene (NCBI RefSeq Gene ID: 4976) comprises 34 exons (30 of which are coding) spanning over 90 kb of genomic DNA. Alternative splicing yields ten distinct isoforms encoding polypeptides of 836 to 1015 amino acids. The resulting OPA1 protein is a dynamin-related GTPase essential for mitochondrial inner membrane organization.

ADOA research relies on an expert-curated *OPA1* database, the Leiden Open Variation Database (LOVD) [5, 6], which encompasses 1,397 genetic variants from 1,687 individuals alongside their clinical classifications. To date, the LOVD lists 746 unique *OPA1* variants, 78% of which are classified as pathogenic (Table I). While the majority (67.4%) are single-nucleotide substitutions, 16% are estimated to impair mRNA splicing. With its 34 exons, dominant inheritance, and expertcurated clinical data, *OPA1* represents an exceptional paradigm for evaluating predictive models on splice-site mutations. Accordingly, two evaluation sets were constructed: a set of mutations with documented splicing defects, and a set of clinically classified benign substitutions (Fig. 1). Test sets encompassed the extended *OPA1* corpus and an independent multigene panel of spliceogenic mutations to assess model utility across mutation types and cross-gene generalizability. This study then evaluated the accuracy of six distinct models—SpliceAI, AlphaGenome, Borzoi, NTv3 650M, Evo2 40B, and Genos 10B—in various iterations (Table II).

**TABLE I.** Distribution and classification of OPA1 variants.

|  | **Variants**<br>**(% of Total)** | **InDels**<br>**(% of Total)** | **Substitutions**<br>**(% of Total)** |
|---|---|---|---|
| Pathogenic | 582 (78.0%) | 227 (93.4%) | 355 (70.6%) |
| VUS | 78 (10.5%) | 9 (3.7%) | 69 (13.7%) |
| Benign | 86 (11.5%) | 7 (2.9%) | 79 (15.7%) |
| **Total (%)** | 746 (100%) | 243 (32.6%) | 503 (67.4%) |

**TABLE II.**
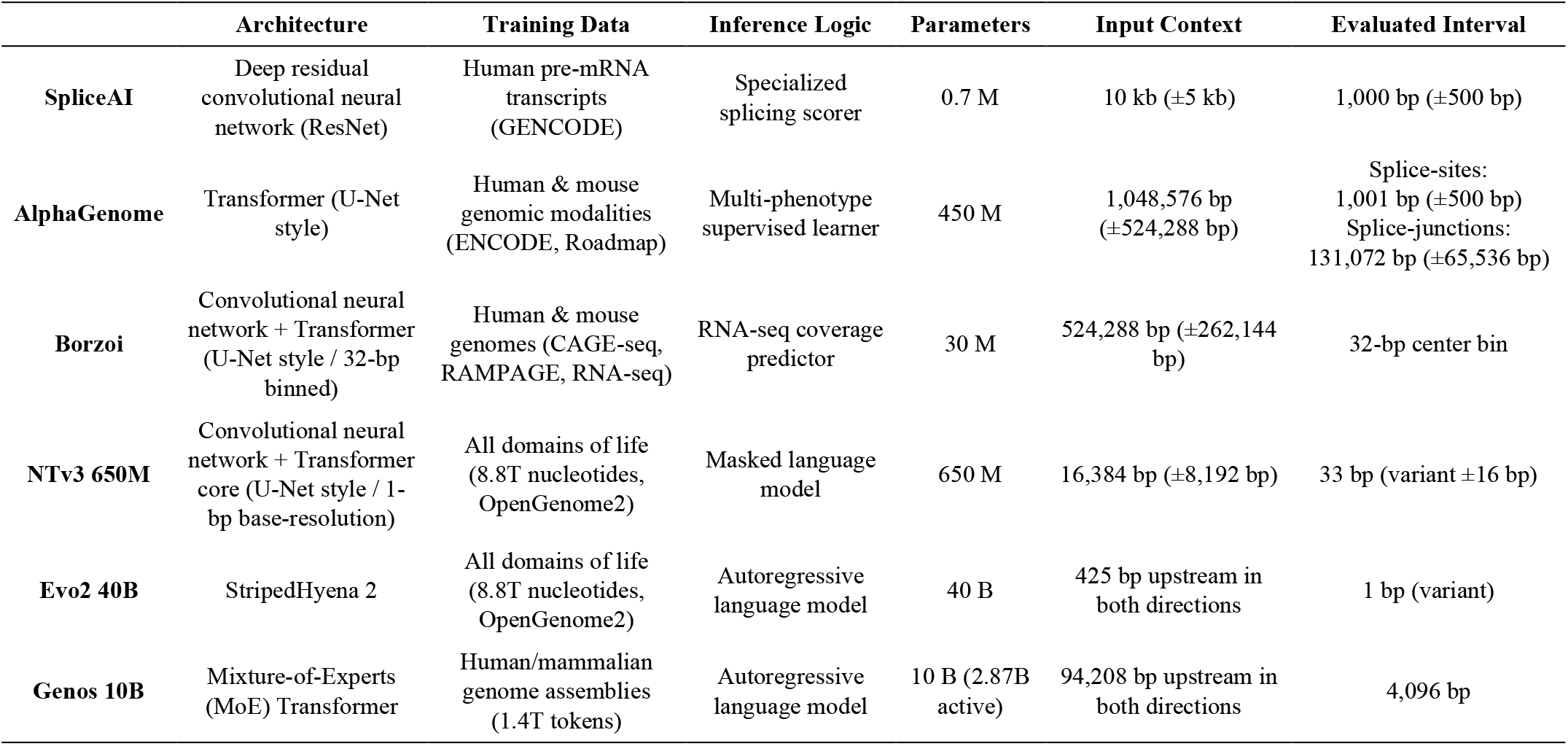
Structural and algorithmic specifications of the different models.

**Fig. 1.**
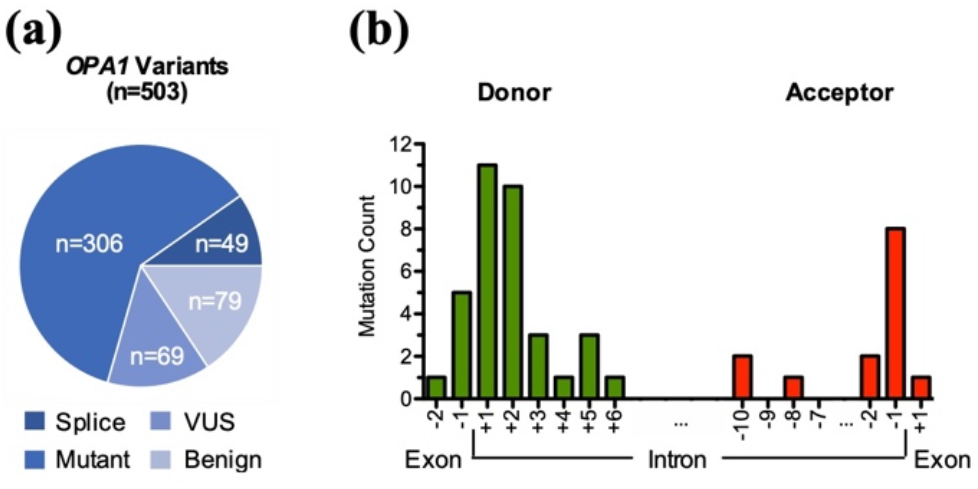
Categorization and positional mapping of *OPA1* variants. **(a)** Partitioning of the 503 unique *OPA1* single-nucleotide substitutions into four distinct sets. **(b)** Positional distribution of the 49 donor (left) and acceptor (right) splice-site mutations relative to exon–intron boundaries.

SpliceAI served as the study’s baseline. The tool utilizes a specialized dilated deep residual convolutional neural network (ResNet) trained on GENCODE human reference transcripts to predict splice sites [7]. AlphaGenome is a polyfunctional encoder-based transformer that leverages attention mechanisms to capture long-range genomic dependencies across distinct molecular phenotypes, such as expression levels and chromatin states [8]. This study evaluated individual AlphaGenome splicing-track outputs alongside the recommended “merged splicing” composite score. Borzoi, by contrast, is a hybrid sequence-to-function architecture trained on next-generation sequencing reads to predict cell- and tissue-specific RNA-seq coverage [9]. This model combines convolutional layers with transformer blocks to capture long-range interactions. Notably, it operates at a spatial resolution that upsamples to a 32-bp output, whereas the other evaluated models provide single-nucleotide resolution.

In contrast to these task-specific, supervised architectures, the three self-supervised genomic foundation models leverage semantic features to enable zero-shot predictions of diverse functional effects. This study calculated relative probabilities for reference and alternative alleles using variant-site logit extraction conditioned on optimized genomic contexts.

Nucleotide Transformer v3 is a 650M-parameter encoder-based model (NTv3 650M) trained on the OpenGenome2 atlas—a curated dataset of 8.8 trillion non-redundant nucleotides spanning all domains of life [10]. Utilizing a bidirectional transformer architecture, NTv3 650M analyzes variants by simultaneously considering both upstream and downstream contexts—akin to a reader filling in a missing word in the middle of a sentence.

Conversely, Evo2 40B and Genos 10B are causal (autoregressive) models that process sequences linearly, predicting each subsequent nucleotide solely from preceding context—much like a large language model anticipating the next word in a sentence. Evo2 40B—a much larger 40B-parameter model than NTv3 650M—was also trained on OpenGenome2 but utilizes hybrid attention-convolution mechanisms (StripedHyena 2) to capture long-range dependencies [11]. By contrast, the human-centric, 10B-parameter Genos 10B model was trained on 636 *de novo* genome assemblies from the Human Pangenome Reference Consortium and Human Genome Structural Variation Consortium, supplemented in version 2 by East Asian and mammalian RefSeq genomes [12]. Genos 10B utilizes a sparse Mixture-of-Experts (MoE) architecture with 2.87B active parameters per token while maintaining a 1-bp resolution.

Collectively, these six models represent a diverse spectrum of architectures, training paradigms, and inference modalities (Table II). This study evaluated what splicing information each model captured and how to effectively retrieve it to maximize clinical utility. Accordingly, hyperparameters were optimized for accuracy and specificity for classifying *OPA1* splice-variants prior to model benchmarking, which revealed mechanistic insights into the representation boundaries of the different architectures.

## II. Methods

All genomic coordinates and annotations are based on the GRCh38 (hg38) genome assembly. All *OPA1* annotations reference transcript variant 1 (NM_015560.2; NP_056375.2).

### A. Dataset selection and curation

A total of 749 unique *OPA1* DNA variants together with their clinical classifications following the American College of Medical Genetics and Genomics and the Association for Molecular Pathology guidelines were retrieved on April 15, 2026 from the Leiden Open Variation Database (LOVD; databases.lovd.nl/shared/genes/OPA1). Following manual curation, three entries were excluded due to recording errors or incorrect annotations, resulting in a dataset of 746 variants.

Clinical significance was stratified into three categories:

- pathogenic/likely pathogenic,
- benign/likely benign, and
- variants of unknown significance (VUS).

For variants with multiple reports, the most severe clinical classification was retained.

From the 503 unique single-nucleotide substitutions, two evaluation sets were established:

- a splicing set comprising 49 mutations annotated with the “consequence: splicing defect”, and
- a benign set comprising 79 benign substitutions.

The remaining 375 *OPA1* substitutions were divided into two additional test panels:

- an extended *OPA1* set of 306 pathogenic variants spanning diverse mutation types, and
- a set of 69 VUS.

To complement the *OPA1* data, a multi-gene test set was constructed from 76 functionally validated, spliceogenic substitutions within the canonical splice sites of 65 genes [13, 14]. Benign controls for these 65 genes were retrieved from ClinVar, filtering for high-confidence single-nucleotide intronic variants reviewed by an expert panel or supported by multiple submitters without conflicting interpretations. To ensure high confidence, “likely benign” substitutions were excluded if the target gene was already represented by a “benign” variant. This filtering yielded a final set of 107 high-confidence benign substitutions spanning 46 of the 65 genes.

### B. Model optimization

Clinical classifications of the *OPA1* evaluation set served as the reference standard. Decision thresholds for each model were determined via a 10,000-parameter one-dimensional grid search, optimizing accuracy and specificity on these variants before expanding analysis to the broader corpus of *OPA1* mutations, VUS, and the multi-gene panels. Performance was further characterized using receiver operating characteristic (ROC-AUC) and precision-recall (PR-AUC) curves.

Primary evaluation metrics were defined as percentages according to Equations 1–5:

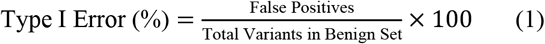

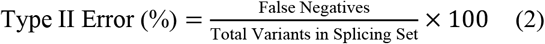

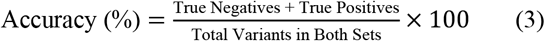

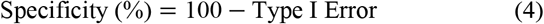

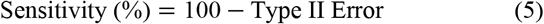

As a secondary metric, positive predictive value (PPV) was calculated according to Equation 6:

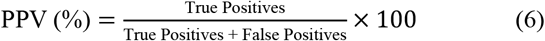

### C. SpliceAI

Delta scores for each variant were retrieved without the masking option from the Broad Institute API (version v1.3.1; github.com/broadinstitute/SpliceAI-lookup) using a 1,000-bp lookup interval centered on the variant position (±500 bp) [7]. A single maximum delta score threshold across the model’s four prediction metrics—acceptor loss, acceptor gain, donor loss, and donor gain—was established using grid search, and variants with a delta score greater than 0.40 were classified as having a pathogenic splicing effect.

### D. lphaGenome

AlphaGenome was accessed via the Google Cloud API through the Google DeepMind Research Access program (deepmind.google.com/science/alphagenome) [8]. The variant classification thresholds for the 4 analyzed outputs were determined via grid search optimization.

Composite splicing scores were computed following AlphaGenome’s splicing-variant-scoring Jupyter notebook for batch processing (colabs/v1/splicing_variant_scoring.ipynb). For each variant, a 1,048,576-bp interval centered on the variant was scored against the reference using the three recommended splicing variant scorers: `SPLICE_SITES`, `SPLICE_SITE_USAGE`, and `SPLICE_JUNCTIONS`. The final composite score was calculated as the weighted sum of maximum predictions across the tracks for splice-sites (*weight* = 1.0), splice-site usage (*weight* = 1.0), and splice-junctions (*weight* = 0.2). In addition, splice-site usage scores were evaluated separately. Variants were classified as having a pathogenic effect if their composite score was greater than or equal to 2.60 or their splice-site usage score greater than or equal to 0.93.

Splice-site effects were additionally evaluated with a custom scoring routine built on the `predict_variant` method (`SPLICE_SITES` output). For each variant, a 1,048,576-bp interval centered on its position was submitted to the model, and splice-site probabilities were obtained for the reference and alternative alleles across the four strand-specific tracks (donor and acceptor, plus and minus strand). Differences in class assignment probabilities were computed for each position within a 1,001-bp window centered on the variant (±500 bp, and the largest decrease (loss) and largest increase (gain) recorded per track. The maximum absolute difference across the four tracks was utilized as the final prediction score, and variants with a score greater than or equal to 0.32 were classified as pathogenic.

Furthermore, splice-junction effects were evaluated with a custom routine built on the `predict_variant` method (`SPLICE_JUNCTIONS` output) using a 131,072-bp interval centered on the variant (±65,536 bp). The log fold change (LFC) between alternate (Alt) and reference (Ref) alleles was calculated across tissue tracks as: LFC = ln(*Alt* + *ε*) − ln(*Ref* + *ε*), with *ε* = 10^−8^. For each junction, the tissue track with the largest absolute log-fold change value was retained and utilized as the final prediction score. Variants with a log-fold change score of greater than or equal to 2.63 were classified as pathogenic.

### E. Borzoi/Flashzoi

Borzoi variant effect predictions [9] were generated via the Flashzoi implementation (github.com/johahi/borzoi-pytorch) [15]. Inference was performed using an ensemble of four pretrained replicate weights (`johahi/borzoi-replicate-0` through -3, retrieved from Hugging Face) in an Ubuntu 24.04 environment on a Virtual Machine equipped with an NVIDIA RTX A5000 (24GB VRAM), utilizing Python 3.12.3, PyTorch 2.8.0, and CUDA 12.8.93.

Borzoi predicts RNA-seq coverage across 7,611 human cell and tissue types. Here, analysis was restricted to 22 plusstrand tracks from the ENCODE project [16] encompassing primary human epithelial, endothelial, mesenchymal, and neural cells from the cardiovascular, respiratory, digestive, nervous, musculoskeletal, renal, reproductive, and integumentary systems [17]. For each variant, the 524,288-bp genomic context window was anchored such that the variant position was centered within the central bin. For both the reference and alternative alleles, RNA-seq read coverage predictions were generated using both forward and reverse-complement sequence orientations, averaged to mitigate strand bias. These values were logarithmically transformed (ln(*Ref*) + 10^!”^ and ln(*Alt*) + 10^!”^, respectively) and then averaged across the four model replicates. Subsequently, values were averaged across the tissue tracks to derive a single consolidated reference and alternative score for each variant. A consensus log fold change was then calculated as LFC = ln(*Alt*) − ln(*Ref*). Because *OPA1* log-fold change scores were not normally distributed, an asymmetric thresholding strategy was implemented (D’Agostino & Pearson test: 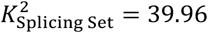 and 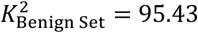 both *p* < 0.0001). The lower threshold (loss) was determined via grid search optimization, whereas the upper boundary (gain) was set at the maximum benign log-fold change score. Variants were classified as having a pathogenic effect if they met either threshold of LFC_lower_ ≤ −0.14 or LFC_upper_ > +0.24 at 32-bp resolution, and LFC_lower_ ≤ −0.10 or LFC_upper_ > +0.19 at 128-bp resolution.

### F. NTv3 650M Pre/Post

Zero-shot causal inference was performed using Nucleotide Transformer NTv3 650M (`InstaDeepAI/NTv3_650M_pre` and `InstaDeepAI/NTv3_650M_post`, retrieved from Hugging Face) [10] in an Ubuntu environment on a Virtual Machine equipped with an NVIDIA A40 (48GB VRAM). The software stack comprised Python 3.11.10, PyTorch 2.4.1, and CUDA 12.4.131, with model loading orchestrated via Hugging Face `transformers` (v4.57.3) and `accelerate` (v1.13.0). Sequences were processed with a batch size of 12 using the model’s native tokenization.

The optimal context size of 16,384 bp (±8,192 bp) was determined empirically using the `NTv3_650M_post` model restricted to human species embeddings via the model call `.encode_species([‘human’])`. Log-likelihoods (LL) for reference and alternative alleles were computed by applying a log-softmax transformation to the model’s output logits and, where indicated, by summing them (∑LL) over a 33-bp interval centered on the variant position (±16 bp). Log-likelihood ratios (LLR = LL(*Alt*) − LL(*Ref*) or LLR = ∑LL(*Alt*) − ∑LL(*Ref*)) were separately calculated for both forward (Fwd) and reverse-complement (RC) strand inferences, and averaged as denoted. Thresholds were established via grid search optimization to classify variants as pathogenic if they met either directional threshold: LLR_304_ ≥ +15.58 or LLR_56_ ≥ +6.56 for NTv3 650M Post, and LLR_304_ ≤ −2.29 or LLR_56_ ≥ +1.26 for NTv3 650M Pre.

### G. Evo2 40B

Autoregressive masked inference was performed using Evo2 40B (version evo2-40b:2.0.0) available from the NVIDIA API catalog via NVIDIA NIM Microservices (build.nvidia.com/arc/evo2-40b) [11]. Inference was executed in both the forward and reverse-complement strand directions using a 425-bp genomic prefix window immediately upstream of each variant position. This prefix was prepended with a human taxonomic identifier (Hu. Id.) to condition the model where indicated. Next-token probabilities were extracted for the reference and alternative alleles applying a log-softmax transformation to the model’s output logits of the hidden state immediately preceding the variant position. Variant effects were quantified using the log-likelihood ratio (LLR = LL(*Alt*) − LL(*Ref*)), calculated separately for both strand orientations. Strand-dependent classification thresholds were established via grid search optimization. Variants were classified as pathogenic if they met either directional threshold: LLR_Fwd_ ≤ −5.34 or LLR_Rc_ ≤ −7.03 without human conditioning, and LLR_Fwd_ ≤ −5.69 or LLR_Rc_ ≤ −5.55 with human conditioning.

### H. Genos 10B

Zero-shot causal inference was performed using the Genos 10B or Genos 10B v2 model (`ZhejiangLab/Genos-10B`, or `ZhejiangLab/Genos-10B-v2`, retrieved from Hugging Face) [12] in an Ubuntu 24.04 environment on a Virtual Machine equipped with an NVIDIA A100 (80GB VRAM) or an NVIDIA RTX Pro 6000 Blackwell (96GB VRAM). The software stack consisted of Python 3.12.3, PyTorch 2.8.0, and CUDA 12.8.93, with model loading orchestrated via Hugging Face `transformers` (v4.46.3) and `accelerate` (v1.14.0). Sequences were evaluated at batch size of 1 (A100) or 2 (RTX Pro 6000 Blackwell) using character-level tokenization and left-padding to maintain right-aligned causal context windows. Inference used FlashAttention-2 in native `torch.bfloat16` precision.

The model processed forward and reverse-complement strand directions independently using an equivalent 94,208-bp genomic window immediately upstream of each variant position. To compute log-likelihoods for reference and alternative alleles, a log-softmax transformation was applied to the model’s output logits. These values were then summed from the variant token index through the 4,096-bp downstream readout interval. Log-likelihood ratios (LLR = ∑LL(*Alt*) − ∑LL(*Ref*)) were calculated separately for the forward and reverse-complement strand inferences, and classification thresholds were established by grid search optimization for each orientation. Variants were flagged as pathogenic if they met either directional threshold: LLR_304_ ≤ −60.57 or LLR_56_ ≤ −98.94.

In addition, the custom Genos 10B model finetuned on ClinVar variants (Genos ClinVar) was evaluated separately. Genos ClinVar-based probability scores were retrieved from the Zhejiang Lab API, accessible through the DCS Cloud catalog (www.dcs.cloud/models/#/genos; source code available at github.com/zhejianglab/Genos). The classification threshold was determined via grid search optimization, and variants with a pathogenic probability of greater than or equal to 0.63 were classified as pathogenic.

### I. Statistical analysis

All statistical analyses, including the coefficient of determination (*R*^2^) for linear regressions, the Friedman test for paired non-parametric data followed by Dunn’s multiple comparison post-hoc test (with SpliceAI serving as the baseline), and D’Agostino & Pearson *K*^2^ normality testing, were performed using Prism 5 (GraphPad Software). Statistical significance was defined *a priori* at α = 0.05. All data are presented as mean ± standard deviation (S.D.) unless indicated otherwise. Normalized global entropy across Borzoi coverage tracks, high-dimensional data processing, model evaluations (ROC-AUC and PR-AUC), and one-dimensional grid searches were computed via custom Python scripts in Google Colaboratory.

## III. Results

Two distinct evaluation sets were established from a total of 503 *OPA1* single-nucleotide substitutions retrieved from LOVD. The splicing set comprised 49 mutations within splicing regions. The benign set—a more genomically distributed control— encompassed 79 substitutions across splicing regions, deep intronic sites, and both synonymous and nonsynonymous exonic positions (Table I; Fig. 1). Hyperparameters of each model were optimized to maximize accuracy and specificity on these evaluation sets before assessing model performance on the test sets, which encompassed the extended *OPA1* corpus and an independent multigene panel of spliceogenic mutations.

### A. Cross-model comparison

Model performance across the *OPA1* evaluation sets was largely comparable. Aside from minor variations in accuracy— from 86.7% to 96.1%—specialized predictors and genomic foundation models performed similarly, with the AlphaGenome iterations achieving top marks on par with SpliceAI (Table III). Likewise, Evo2 40B demonstrated comparable sensitivity (93.9%) to the specialized models, albeit with a slightly higher type I error rate. In contrast, Genos 10B achieved an accuracy of 77.3%, significantly underperforming relative to all other architectures, primarily due to a low sensitivity of 49.0%. The finetuned Genos ClinVar model showed a further reduction in these metrics. Aside from the underperformance of the Genos 10B iterations, differences among the models remained within statistical noise (Table III; Fig. 2).

**Fig. 2.**
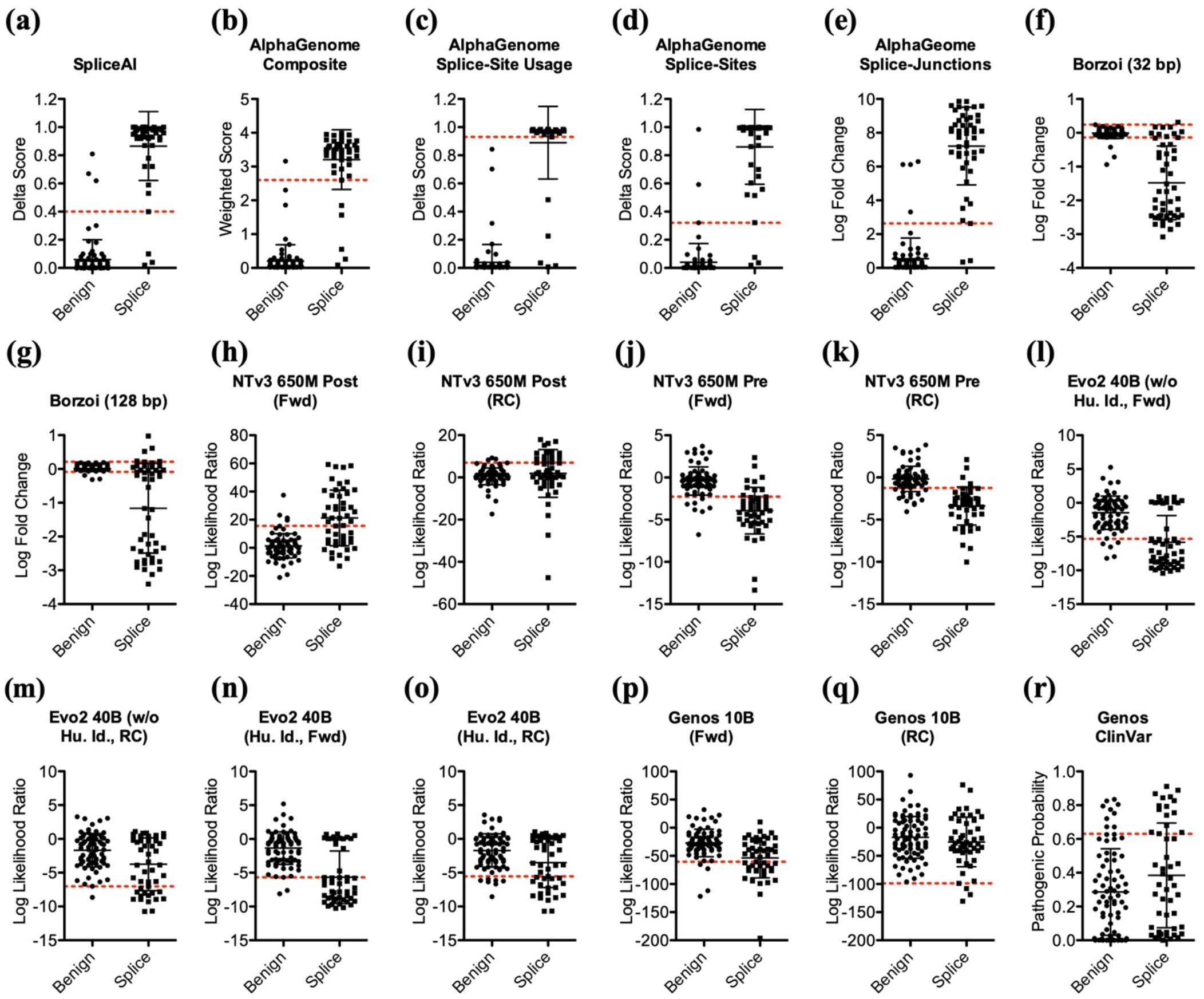
OPA1 variant classification across model architectures. Variant prediction scores for the evaluation set across evaluated models, with horizontal red dashed lines indicating optimal classification thresholds. **(a–e)** Splicing-focused model scores, including SpliceAI delta scores **(a)**, AlphaGenome composite scores **(b)**, splice-site usage predictions **(c)**, splice-site scores **(d)**, and splice-junction log-fold change values **(e). (f, g)** Borzoi log-fold change values at 32-bp **(f)** and 128-bp **(g)** bin sizes, where upper and lower thresholds denote predicted splice-site gains and losses, respectively. **(h–q)** Genomic foundation models log-likelihood ratios (LLRs) across forward (Fwd) and reverse-complement (RC) strands for post-trained NTv3 650M Post **(h, i)**, pretrained NTv3 650M Pre **(j, k)**, Evo2 40B without human taxonomic identifier (Hu. Id.) **(l, m)**, Evo2 40B with human taxonomic identifier **(n, o)**, and Genos 10B **(p, q)**. Variants were classified as pathogenic if scores exceeded either directional threshold. **(r)** Genos ClinVar pathogenicity scores.

**TABLE III.** Model performance on the *OPA1* corpus.

|  | **Evaluation Sets** |  | ***OPA1* Benign (n=79)** |  | ***OPA1* Splicing (n=49)** |  | **Extended *OPA1* Corpus (n=306+69)** |  |
|---|---|---|---|---|---|---|---|---|
|  | **Accuracy** | **PPV** | **Type I<br>Error** | **Specificity** | **Sensitivity** | **Type II<br>Error** | **Positive<br>Mutations** | **Positive<br>VUS** |
| **SpliceAI** | 95.3% | 93.9% | 3 (3.8%) | 76 (96.2%) | 46 (93.9%) | 3 (6.1%) | 75 (24.5%) | 6 (8.7%) |
| **AlphaGenome<br>Composite** | 95.3% | 97.8% | 1 (1.3%) | 78 (98.7%) | 44 (89.8%) | 5 (10.2%) | 69 (22.5%) | 3 (4.3%) |
| **AlphaGenome<br>Splice-Site Usage** | 96.1% | 100.0% | 0 (0.0%) | 79 (100%) | 44 (89.8%) | 5 (10.2%) | 60 (19.6%) | 2 (2.9%) |
| **AlphaGenome<br>Splice-Sites** | 96.1% | 95.8% | 2 (2.5%) | 77 (97.5%) | 46 (93.9%) | 3 (6.1%) | 74 (24.2%) | 3 (4.3%) |
| **AlphaGenome<br>Splice-Junctions** | 95.3% | 92.2% | 4 (5.1%) | 75 (94.9%) | 47 (95.9%) | 2 (4.1%) | 88 (28.8%) | 8 (11.6%) |
| **Borzoi (32-bp)** | 91.4% | 93.2% | 3 (3.8%) | 76 (96.2%) | 41 (83.7%) | 8 (16.3%) | 86 (28.1%) | 7 (10.1%) |
| **Borzoi (128-bp)** | 89.1% | 92.9% | 3 (3.8%) | 76 (96.2%) | 39 (79.6%) | 10 (20.4%) | 70 (22.9%) | 13 (18.8%) |
| **NTv3 650M Post** | 89.1% | 88.9% | 5 (6.3%) | 74 (93.7%) | 40 (81.6%) | 9 (18.4%) | 77 (25.2%) | 9 (13.0%) |
| **NTv3 650M Pre** | 86.7% | 77.6% | 13 (16.5%) | 66 (83.5%) | 45 (91.8%) | 4 (8.2%) | 212 (69.3%) | 23 (33.3%) |
| **Evo2 40B<br>(w/o Hu. Id.)** | 93.0% | 88.5% | 6 (7.6%) | 73 (92.4%) | 46 (93.9%) | 3 (6.1%) | 280 (91.5%) | 46 (66.7%) |
| **Evo2 40B<br>(Hu. Id.)** | 92.2% | 86.8% | 7 (8.9%) | 72 (91.1%) | 46 (93.9%) | 3 (6.1%) | 276 (90.2%) | 47 (68.1%) |
| **Genos 10B** | 77.3% | 85.7% | 4 (5.1%) | 75 (94.9%) | 24 (49.0%) | 25 (51.0%) | 92 (30.1%) | 23 (33.3%) |
| **Genos<br>ClinVar** | 66.4% | 62.5% | 9 (11.4%) | 70 (88.6%) | 15 (30.6%) | 34 (69.4%) | 104 (34.0%) | 25 (36.2%) |

### B. Model tuning

An assessment of SpliceAI’s delta scores revealed that individual variants can generate concurrent gains and losses at both acceptor and donor sites, indicating localized shifts in splice-site selection. Accordingly, a single maximum delta score threshold across the model’s four prediction metrics was established to maximize classification accuracy (Fig. 2a; Supplementary Fig. S1). AlphaGenome’s recommended method to assess variants for their splicing impact integrates three distinct outputs from the model’s splice-site, splice-site usage, and splice-junction modules. On the evaluation sets, the composite and the individual modules achieved the highest accuracy and lowest type I error rates (Table III). Splice-site usage predicts how efficiently or frequently particular splice sites are utilized relative to one another. The splice-site module quantifies changes in class assignment probabilities for gains and losses at acceptor and donor sites, thereby capturing local shifts in splice-site probabilities similar to SpliceAI. Notably, only the forward strand orientation was informative, while the reverse-complement strand did not add any predictive value. Meanwhile, the splice-junction module computes the inner product between processed donor and acceptor embeddings, yielding high-scoring junctions that were often distal (>100 bp) in cases where local scores fail to reach significance (Figs. 2b–2e; Supplementary Fig. S1).

Borzoi, the third supervised model, predicts RNA-seq coverage across 7,611 distinct tracks for a 524,288-bp sequence interval with a 32-bp binned resolution. To account for OPA1’s ubiquitous expression, center-bin coverage across 22 primary human epithelial, endothelial, mesenchymal, and neural cell tracks was averaged to derive a consensus log-fold change score [17]. Because these scores were non-normally distributed, an asymmetric thresholding strategy was implemented to isolate coverage losses from gains. This approach achieved accuracies of 91.4% at a 32-bp bin resolution and 89.1% at 128-bp (Table III; Figs. 2f and 2g; Supplementary Fig. S2).

Transitioning to the genomic foundation models. For the bidirectional NTv3 650M Transformer—encompassing both pretrained and species-post-trained iterations—initial single-nucleotide inference suffered from high type II error rates regardless of the surrounding genomic context. However, accounting for local vicinity by aggregating log-likelihood ratios across a 33-bp interval considerably improved accuracy. With 16,384 bp context, the post-trained model achieved 89.1% accuracy—counterintuitively for a bidirectional architecture—by combining forward and reverse-complement strand inferences via a logical OR operation. In comparison, the pretrained model yielded 86.7% accuracy by this approach, whereas other configurations underperformed (Table III; Figs. 2h–2k; Supplementary Table I; Supplementary Fig. S3).

Evo2 40B, a hybrid attention-convolution model, excelled at predicting variant effects at single-base resolution with a minimal upstream context of just 425 bp. Asymmetry inherent to the model’s causal architecture necessitated a logical OR operation combing forward and reverse-complement strand inferences— yielding 93.0% accuracy and indicating distinct upstream versus downstream context dependencies across variants. Notably, human preconditioning increased type I errors, while expanding the context window degraded performance. (Table III; Figs. 2l– 2o; Supplementary Table II; Supplementary Fig. S4).

Genos 10B also benefited from OR gating forward and reverse-complement strand inferences. However, the MoE architecture required an expansive 94,208-bp upstream context and downstream logit aggregation across 4,096 bp to reach 77.3% accuracy. Maintaining at least bfloat16 precision was thereby critical to resolve micro-probability shifts across this 4,096-logit vector. Furthermore, migrating the workload across different hardware architectures degraded predictive accuracy, suggesting that platform-specific kernel reductions or accumulator behaviors introduce confounding numerical noise. At the same time, adjusting evaluation windows shifted the balance of type I and type II errors without altering overall accuracy, indicating a performance ceiling. Nonetheless, this optimized Genos 10B configuration outperformed both the finetuned ClinVar model and the extended-species v2 iteration (Table III; Figs. 2p–2r; Supplementary Table III; Supplementary Fig. S5).

### C. Classification errors

At the variant level, c.2708-10C>G (OPA1_000081) and p.Val933Phe (c.2797G>T; OPA1_000460) were the sole exceptions in the splicing set, whereas all other mutations were detected by at least one specialized model. Because both variants were first documented in patients with compound heterozygous mutations [18, 19], they may represent hypomorphic/conditional alleles with subtle functional impacts, clinical ambiguity, or shared algorithmic false negatives. While the c.2708-10C>G substitution was uniquely flagged by NTv3 650M Post, the p.Val933Phe mutation was classified as pathogenic by Evo2 40B and Genos 10B. Meanwhile, false positives within the benign set were scattered stochastically across models (Supplementary Table IV).

### D. Positional variant classification across the OPA1 corpus

For the remaining *OPA1* substitutions within the LOVD corpus, Evo2 40B exhibited a broad detection rate regardless of mutation type, classifying 90.2% of the 306 mutations as pathogenic with human preconditioning and 91.5% without. Similarly, Evo2 40B predicted 66.7% (with preconditioning) and 68.1% (without preconditioning) of the 69 VUS as pathogenic.

By contrast, human species-post-training produced a marked divergence in the NTv3 650M architecture: while the base NTv3 650M Pre model detected 69.3% of *OPA1* mutations, NTv3 650M Post flagged only 25.2% of them, mirroring the sensitivities seen with the supervised models and Genos 10B. A similar shift occurred among the VUS, where positive prediction rates fell from 33.3% (NTv3 650M Pre) to 13.0% (NTv3 650M Post), again mirroring the supervised models (Table III; Fig. 3).

**Fig. 3.**
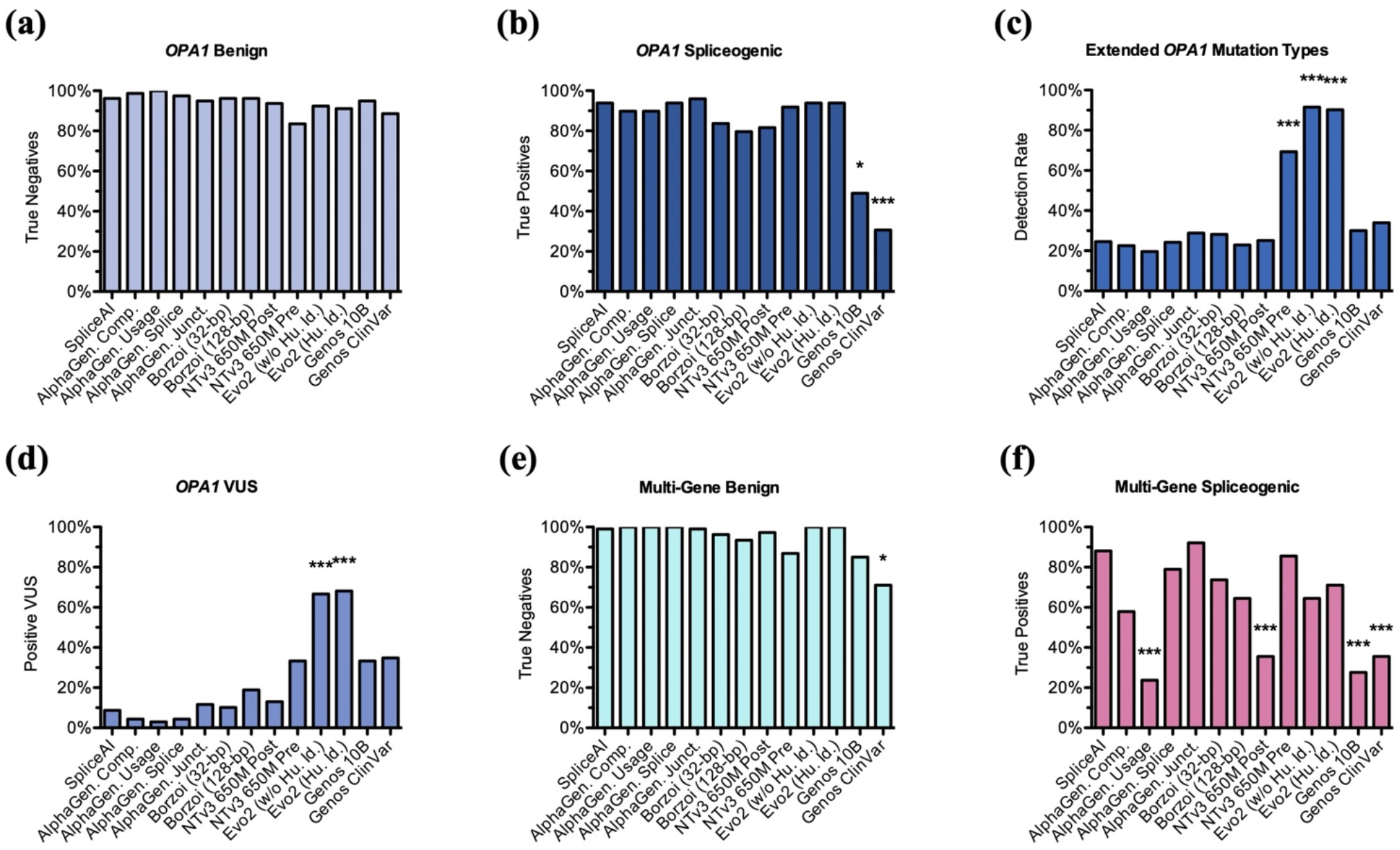
Variant classification across model architectures and cohorts. **(a)** Specificity (true negative rate) across the *OPA1* benign set and **(b)** sensitivity (true positive rate) across the *OPA1* splicing set. **(c)** Detection rates across the extended *OPA1* corpus encompassing various mutation types. **(d)** Positive prediction rate across *OPA1* variants of uncertain significance (VUS). **(e)** Specificity and **(f)** sensitivity evaluated across the multi-gene cohort. Statistical significance relative to SpliceAI was determined using a Friedman test followed by Dunn’s multiple comparison test (\**p*<0.05; \*\**p*<0.01; \*\*\**p*<0.001).

Nonetheless, only the specialists SpliceAI, AlphaGenome, and Borzoi, alongside NTv3 650M Post, identified two deep intronic mutations that give rise to cryptic splice sites: c.625-853T>G (OPA1_000717) and c.678+180A>G (OPA1_000718) [3]. Notably, the pretrained NTv3 650M model failed to identify these cryptic splice sites, revealing a distinct shift in representation boundaries between the model’s pre- and post-trained iterations (Supplementary Table IV).

### E. Positional consistencies

Most of the 45 variant positions with multiple alleles showed consensus across architectures regardless of the nucleotide exchange. For instance, the three point mutations c.1516+1G>T (OPA1_000032), c.1516+1G>A (OPA1_000701), and c.1516+1G>C (OPA1_000702) were recognized near-uniformly as pathogenic, demonstrating that transitions and transversions at this position were consistently captured across models. Also the classification of the mutations c.2013G>A (OPA1_000155) and c.2013G>C (OPA1_000647) was largely consistent among the architectures, as was the classification of the adjacent variants c.2013+1G>T (OPA1_000201), c.2013+1G>A (OPA1_000118), and c.2013+1G>C (OPA1_000299).

This pattern highlights a broader trend in how the different architectures graded positions with multiple alternative alleles. While Genos 10B classified such positions divergently in 19 of 45 instances, the other models maintained higher positional consistency, with discrepancies ranging from 1 to 10 instances.

Beyond general consistency, looking at specific multi-allelic sites revealed nuanced differences in model sensitivity. For instance, only NTv3 650M and Evo2 40B configured with human preconditioning (but not without) accurately differentiated the nonsense mutation c.1096C>T (OPA1_000001) from the benign synonymous variant c.1096C>A (OPA1_000053). Furthermore, the well-characterized nonsynonymous mutations p.Arg445His (c.1334G>A; OPA1_000104) and p.Arg445Leu (c.1334G>T; OPA1_000572) within *OPA1’s* GTPase domain were identified exclusively by Evo2 40B, NTv3 650M Pre, and the finetuned Genos ClinVar model (Supplementary Table V).

### F. Multi-gene benchmarking

To assess generalizability, architectures were benchmarked against an independent 65-gene dataset comprising 107 benign substitutions and 76 confirmed spliceogenic mutations [13, 14]. Most architectures demonstrated comparable performance on this test set, with SpliceAI as well as the AlphaGenome splice-site and splice-junction modules achieving over 90% accuracy. Except for Genos 10B, all models maintained high specificity, whereas sensitivity was generally lower. Notably, AlphaGenome splice-site usage, NTv3 650M Post, and Genos 10B showed significantly lower detection rates. Taken together, the near-perfect specificity of SpliceAI, AlphaGenome, and Evo2 40B—at the cost of slightly reduced sensitivity—demonstrates that *OPA1*-derived thresholds are rather conservative (Table IV; Fig. 3; Supplementary Table VI).

**TABLE IV.** Model performance on the multi-gene corpus.

|  | ***Multi-Gene* Test Sets** |  | ***Multi-Gene* Benign (n=107)** |  | ***Multi-Gene* Splicing (n=76)** |  |
|---|---|---|---|---|---|---|
|  | **Accuracy** | **PPV** | **Type I Error** | **Specificity** | **Sensitivity** | **Type II Error** |
| **SpliceAI** | 94.5% | 98.5% | 1 (0.9%) | 106 (99.1%) | 67 (88.2%) | 9 (11.8%) |
| **AlphaGenome<br>Composite** | 82.5% | 100.0% | 0 (0.0 %) | 107 (100%) | 44 (57.9%) | 32 (42.1%) |
| **AlphaGenome<br>Splice-Site Usage** | 68.3% | 100.0% | 0 (0.0 %) | 107 (100%) | 18 (23.7%) | 58 (76.3%) |
| **AlphaGenome<br>Splice-Sites** | 91.3% | 100.0% | 0 (0.0 %) | 107 (100%) | 60 (78.9%) | 16 (21.1%) |
| **AlphaGenome<br>Splice-Junctions** | 96.2% | 98.6% | 1 (0.9%) | 106 (99.1%) | 70 (92.1%) | 6 (7.9%) |
| **Borzoi (32-bp)** | 86.9% | 93.3% | 4 (3.7%) | 103 (96.3%) | 56 (73.7%) | 20 (26.3%) |
| **Borzoi (128-bp)** | 80.9% | 87.5% | 7 (6.5%) | 100 (93.5%) | 49 (64.5%) | 27 (35.5%) |
| **NTv3 650M Post** | 71.6% | 90.0% | 3 (2.8%) | 104 (97.2%) | 27 (35.5%) | 49 (64.5%) |
| **NTv3 650M Pre** | 86.3% | 82.3% | 14 (13.1%) | 93 (86.9%) | 65 (85.5%) | 11 (14.5%) |
| **Evo2 40B<br>(w/o Hu. Id.)** | 85.2% | 100% | 0 (0.0 %) | 107 (100%) | 49 (64.5%) | 27 (35.5%) |
| **Evo2 40B<br>(Hu. Id.)** | 88.0% | 100% | 0 (0.0 %) | 107 (100%) | 54 (71.1%) | 22 (28.9%) |
| **Genos 10B** | 61.7% | 56.8% | 16 (15.0%) | 91 (85.0%) | 21 (27.6%) | 55 (72.4%) |
| **Genos<br>ClinVar** | 56.3% | 46.6% | 31 (28.7%) | 76 (71.0%) | 27 (35.5%) | 49 (64.5%) |

### G. Secondary metric analysis

Threshold selection was optimized to minimize type I errors, resulting in uniformly high positive predictive values across both evaluation and test sets. However, plotting sensitivity on the spliceogenic multi-gene panel against the corpus of extended *OPA1* mutations separated the foundation models from the splicing specialists, as Evo2 40B and NTv3 650M Pre detected significantly more *OPA1* mutations than the other models (Fig. 4). Grid search profiles revealed that SpliceAI and AlphaGenome maintain high classification accuracy across a broad threshold range, whereas other models exhibited narrower optimal windows (Supplementary Fig. S6). Receiver operating characteristic analysis confirmed strong discrimination across task-specific architectures. SpliceAI and AlphaGenome achieved ROC-AUC values exceeding 0.98, while Borzoi reached 0.88. Conversely, generalists yielded lower ROC-AUC scores and exhibited marked strand sensitivity, dropping from 0.73–0.81 on the forward strand (Genos 10B to NTv3 650M Post) to 0.55–0.63 on the reverse-complement strand (Genos 10B to Evo2 40B). NTv3 650M Pre was the exception, maintaining robust performance across both orientations (ROC-AUC Fwd: 0.91; RC: 0.89). Precision-recall analysis confirmed this performance split between task-specific splice predictors and generalist foundation models across the *OPA1* evaluation sets (Supplementary Figs. S7 and S8).

**Fig. 4.**
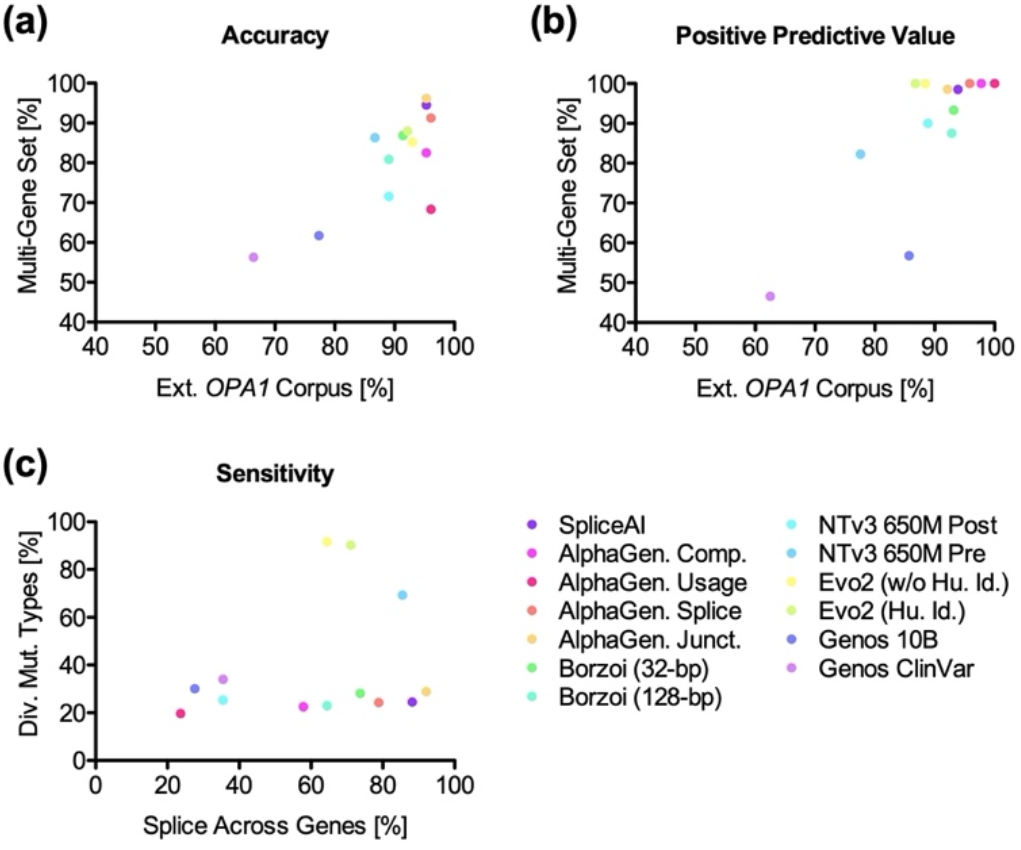
Comparative performance metrics across models. **(a)**Accuracy (%) on the multi-gene dataset vs. the extended *OPA1* corpus. **(b)** Positive predictive value (PPV, %) across datasets, showing high clustering driven by stringent type I error thresholding. **(c)** Sensitivity (%) across diverse *OPA1* mutation types vs. spliceogenic mutations across genes.

## IV. Discussion

The division between task-specific supervised architectures and self-supervised foundation models is blurring. This study focused on classification accuracy as the primary evaluation metric. Although regulatory frameworks prioritize both accuracy and positive predictive value when assessing diagnostic utility, the latter proved uninformative due to threshold selection minimizing type I error rates. Notably, the resulting higher specificity also masked reduced model performance on the multi-gene panel indicating that the *OPA1*-derived classification thresholds are conservative.

Supervised architectures excel on low-noise, task-specific functional data, a paradigm also demonstrated for splicing predictions [20]. Here, SpliceAI’s ResNet architecture and targeted training efficiently mapped variants to localized reconfigurations of splice-site probabilities without generalized semantic modeling. Similarly, Borzoi accurately classified splice-site mutations based strictly on predicted RNA-seq coverage profiles, omitting explicit semantic modeling of gene regulation, transcription initiation, or mRNA maturation. This performance generalized to the multi-gene cohort, confirming that inferred RNA-expression profiles effectively serve as a functional proxy for splicing events.

AlphaGenome similarly achieved high accuracy on splicing tasks; yet, its recommended composite scoring framework underperformed relative to its isolated modules. Specifically, the individual splice-site and splice-junction modules demonstrated superior accuracy and generalizability to the composite framework even surpassing SpliceAI’s accuracy. The relative performance drop of the splice-site usage module on the multi-gene panel, on the other hand, likely stems from its threshold selection routine, which selects the rightmost local peak threshold.

SpliceAI scales linearly with sequence length via dilated convolutions. In contrast, transformer-based models such as AlphaGenome and Borzoi incur quadratic attention scaling, which they mitigate through sequence downsampling to 128-bp resolution bins. Despite this spatial resolution loss, both architectures effectively resolved single-nucleotide splice-site mutations, generalizing across the multi-gene cohort and achieving performance comparable to SpliceAI.

Genomic foundation models inherently demand high-volume data and substantial compute, but extracting high-fidelity splicing representations may depend specifically on evolutionarily diverse sequence data. For example, NTv3 650M and Evo2 40B were both trained on the pan-species OpenGenome2 dataset, yet their allocation of parameter scale, input context, and architectural inductive biases diverged considerably. NTv3 650M’s compact transformer architecture required 16,384 bp of genomic context and logit aggregation across a 33-bp interval to classify *OPA1* variants. Evo2 40B’s massive multi-hybrid architecture, in contrast, predicted *OPA1* variant effects efficiently using a narrow 425-bp context window paired with a single-token logit, challenging the assumption that input context length is the primary driver of splicing fidelity [20]. The divergence between both models points instead to an asymmetry in computational cost distribution, whereby overhead is either frontloaded onto parameter scaling to minimize inference-time context, or backloaded onto deployment through expansive context windows and logit aggregation.

In comparison, Genos 10B—constrained by both a smaller, human-centric training corpus and a sparse MoE architecture optimized for macro-scale regulatory landscapes—required an enormous 94,208-bp context window and logit aggregation across a huge 4,096-bp interval to isolate functional variant effects. This dependency suggests that the model’s capacity to resolve local splicing syntax is limited without diverse evolutionary context. Yet, broadening training distributions across mammalian genomes degraded Genos 10B v2’s performance, pointing to potential routing conflicts within the sparse MoE layers.

Conversely, Evo2 40B’s multi-hybrid architecture preserves single-nucleotide resolution by alternating input-dependent convolutional operators with attention layers. Activating its human-refined configuration shifted the embedding space to correctly differentiate the benign synonymous c.1096C>A variant OPA1_000001 from the nonsense c.1096C>T mutation OPA1_000053 at this same genomic position. Human preconditioning similarly aligned the post-trained NTv3 650M Post model with the specialized architectures, identifying the deep intronic cryptic splice sites OPA1_000717 and OPA1_000718, while missing nonsynonymous mutations such as OPA1_000104. In contrast, the pretrained baseline NTv3 650M Pre more closely mirrored the generalists’ trade-off between identifying nonsynonymous mutations and missing cryptic splice sites.

Convergent accuracies across model categories indicate that diagnostic utility is not inherently restricted by model class, but rather emerges from the intricate interplay of architecture, training paradigm, and inference modality. For example, Evo2 40B reached 88.0% accuracy on the multi-gene test set, surpassing AlphaGenome’s recommended composite score and Borzoi, while approaching SpliceAI’s performance. Comparing AlphaGenome, Borzoi, and NTv3 650M—which all share a U-Net transformer backbone—demonstrates that supervised multi-task conditioning and self-supervised generalist training achieve parity. This convergence underscores machine learning’s “bitter lesson”: scale and uncurated context can compensate for a lack of explicit functional tuning [21].

Two methodological constraints inherent to this study’s design—single-gene threshold derivation and calibration restricted to spliceogenic variants—illuminate two architectural vulnerabilities:

First, single-gene thresholding on *OPA1* exposed clear differences in generalizability across the multi-gene panel. While several architectures transferred reliably, AlphaGenome splice-site usage, NTv3 650M Post, and both Genos 10B iterations suffered significant drops in detection rates. Although part of this reduction reflects a grid search routine that selected conservative thresholds, the performance split suggests that decision boundaries optimized on a single locus may not uniformly generalize across genomic contexts.

Second, restricting threshold derivation to spliceogenic variants limited performance across diverse mutation classes. Consequently, human-centered foundation models like NTv3 650M Post and Genos 10B—alongside task-specific models—underperformed when evaluating the extended corpus of diverse *OPA1* mutation types. This limitation of class-specific thresholding further highlights a structural bottleneck for multi-task architectures, such as AlphaGenome, where the requirement for task-specific thresholding for each distinct mutation type limits their extensibility. Conversely, NTv3 650M Pre and Evo2 40B maintained high sensitivity across mutation types and genes, indicating that locus-specific calibration—rather than mutation-class calibration—drives clinical utility in genomic foundation models. Moreover, it signals that self-supervised generalists will soon surpass task-specific supervised networks in resolving clinical variants.

## V. Conclusion

Evaluating foundation models solely across benchmarks constitutes an insufficiently specified problem setting, widening the disconnect between leader-board metrics and real-world utility. As formalized by Goodhart’s law, optimizing against static evaluation targets encourages dataset artifact exploitation, yielding nominal performance gains that fail to reflect robust generalization.

Moving beyond broad benchmarks, this study probes how generalist architectures internalize RNA splicing syntax at single-nucleotide resolution. By first optimizing on an expert-curated, single-gene locus before extending testing to a multi-gene cohort, this granular approach delineates three core principles governing genomic foundation models:

1. Training paradigms fundamentally dictate performance: extracting splicing syntax requires broad datasets that span evolutionary dimensions.
2. Parameter scale does not guarantee syntax internalization; capacity is only one side of a compute trade-off that shifts overhead between training frontloading and inference backloading.
3. Architecture exerts decisive control over model stability: a sparse MoE model experienced collapse during single-nucleotide variant classification, despite holding state-of-the-art benchmark ranks.

In summary, this work demonstrates the limitations of relying on coarse benchmark metrics to gauge genomic model capabilities, introducing instead a targeted, mechanistically grounded evaluation framework. By establishing that data diversity, compute allocation, and architectural stability supersede raw parameter scale, this study reframes how genomic foundation models should be benchmarked, stress-tested, and built.

## Acknowledgment

During preparation of this manuscript, the author used Claude Sonnet 5 (Anthropic) and Gemini 2.5 Flash (Google Colab) for coding assistance, and Gemini 3.5 Flash (Google) for manuscript editing and proofreading. The author manually reviewed all output and maintains full responsibility for the scientific integrity and final content of this work.

## Software Availability

All source code and instructions are publicly available on GitHub (github.com/marcel-2c8/splicing-model-eval-2026).

