## Supplementary Materials for "Probing Genomic Foundation Models with Splice-Variant Perturbations"

Marcel V. Alavi

SUPPLEMENTARY TABLE I
PERFORMANCE METRICS OF NUCLEOTIDE TRANSFORMER V3 (NTV3) ITERATIONS.

| **NTv3 650M Post** | **Fwd** | **RC** | **Fwd/RC Avg.** | **Fwd OR RC** |
|---|---|---|---|---|
| **Benign Set (n=79)** | | | | |
| False Positives (Type I Error) | 4 (5.1%) | 3 (3.8%) | 7 (8.9%) | 5 (6.3%) |
| True Negatives (Specificity) | 75 (94.9%) | 76 (96.2%) | 72 (91.1%) | 74 (93.7%) |
| **Splicing Set (n=49)** | | | | |
| True Positive (Sensitivity) | 29 (59.2%) | 18 (36.7%) | 38 (77.6%) | 40 (81.6%) |
| False Negative (Type II Error) | 20 (40.8%) | 31 (63.3%) | 11 (22.4%) | 9 (18.4%) |
| **Accuracy** | 81.3% | 73.4% | 85.9% | 89.1% |
| **NTv3 650M Post (w/o Hu. Embed.)** | **Fwd** | **RC** | **Fwd/RC Avg.** | **Fwd OR RC** |
| **Benign Set (n=79)** | | | | |
| False Positives (Type I Error) | 5 (6.3%) | 4 (5.1%) | 9 (11.4%) | 7 (8.9%) |
| True Negatives (Specificity) | 74 (93.7%) | 75 (94.9%) | 70 (88.6%) | 72 (91.1%) |
| **Splicing Set (n=49)** | | | | |
| True Positive (Sensitivity) | 29 (59.2%) | 20 (40.8%) | 38 (77.6%) | 40 (81.6%) |
| False Negative (Type II Error) | 20 (40.8%) | 29 (59.2%) | 11 (22.4%) | 9 (18.4%) |
| **Accuracy** | 80.5% | 74.2% | 84.4% | 87.5% |
| **NTv3 100M Post** | **Fwd** | **RC** | **Fwd/RC Avg.** | **Fwd OR RC** |
| **Benign Set (n=79)** | | | | |
| False Positives (Type I Error) | 1 (1.3%) | 6 (7.6%) | 2 (2.5%) | 7 (8.9%) |
| True Negatives (Specificity) | 78 (98.7%) | 73 (92.4%) | 77 (97.5%) | 72 (91.1%) |
| **Splicing Set (n=49)** | | | | |
| True Positive (Sensitivity) | 14 (28.6%) | 17 (34.7%) | 10 (20.4%) | 26 (53.1%) |
| False Negative (Type II Error) | 35 (71.4%) | 32 (65.3%) | 39 (79.6%) | 23 (46.9%) |
| **Accuracy** | 71.9% | 70.3% | 68.0% | 76.6% |
| **NTv3 650M Pre** | **Fwd** | **RC** | **Fwd/RC Avg.** | **Fwd OR RC** |
| **Benign Set (n=79)** | | | | |
| False Positives (Type I Error) | 7 (8.9%) | 13 (16.5%) | 9 (11.4%) | 13 (16.5%) |
| True Negatives (Specificity) | 72 (91.1%) | 66 (83.5%) | 70 (88.6%) | 66 (83.5%) |
| **Splicing Set (n=49)** | | | | |
| True Positive (Sensitivity) | 40 (81.6%) | 45 (91.8%) | 42 (85.7%) | 45 (91.8%) |
| False Negative (Type II Error) | 9 (18.4%) | 4 (8.2%) | 7 (14.3%) | 4 (8.2%) |
| **Accuracy** | 87.5% | 73.4% | 87.5% | 86.7% |

SUPPLEMENTARY TABLE II
PERFORMANCE METRICS OF EVO2 40B CONFIGURATIONS.

| **Evo2 40B** | **Fwd** | **RC** | **Fwd/RC Avg.** | **Fwd OR RC** |
|---|---|---|---|---|
| **Benign Set (n=79)** | | | | |
| False Positives (Type I Error) | 2 (2.5%) | 7 (8.9%) | 11 (13.9%) | 7 (8.9%) |
| True Negatives (Specificity) | 77 (97.5%) | 72 (91.1%) | 68 (86.1%) | 72 (91.1%) |
| **Splicing Set (n=49)** | | | | |
| True Positive (Sensitivity) | 34 (69.4%) | 19 (38.8%) | 35 (71.4%) | 46 (93.9%) |
| False Negative (Type II Error) | 15 (30.6%) | 30 (61.2%) | 14 (28.6%) | 3 (6.1%) |
| **Accuracy** | **86.7%** | **71.1%** | **80.5%** | **92.2%** |
| **Evo2 40B (w/o Hu. Embed.)** | **Fwd** | **RC** | **Fwd/RC Avg.** | **Fwd OR RC** |
| **Benign Set (n=79)** | | | | |
| False Positives (Type I Error) | 5 (6.3%) | 1 (1.3%) | 12 (15.2%) | 6 (7.6%) |
| True Negatives (Specificity) | 74 (93.7%) | 78 (98.7%) | 67 (84.8%) | 73 (92.4%) |
| **Splicing Set (n=49)** | | | | |
| True Positive (Sensitivity) | 35 (71.4%) | 15 (30.6%) | 38 (77.6%) | 46 (93.9%) |
| False Negative (Type II Error) | 14 (28.6%) | 34 (69.4%) | 11 (22.4%) | 3 (6.1%) |
| **Accuracy** | **85.2%** | **72.7%** | **82.0%** | **93.0%** |
| **Evo2 40B (937 bp/1 bp)** | **Fwd** | **RC** | **Fwd/RC Avg.** | **Fwd OR RC** |
| **Benign Set (n=79)** | | | | |
| False Positives (Type I Error) | 5 (6.3%) | 3 (3.8%) | 13 (16.5%) | 7 (8.9%) |
| True Negatives (Specificity) | 74 (93.7%) | 76 (96.2%) | 66 (83.5%) | 72 (91.1%) |
| **Splicing Set (n=49)** | | | | |
| True Positive (Sensitivity) | 34 (69.4%) | 16 (32.7%) | 38 (77.6%) | 46 (93.9%) |
| False Negative (Type II Error) | 15 (30.6%) | 33 (67.3%) | 11 (22.4%) | 3 (6.1%) |
| **Accuracy** | **84.4%** | **71.9%** | **81.3%** | **92.2%** |

SUPPLEMENTARY TABLE III

COMPARISON OF GENOS 10B PERFORMANCE METRICS ACROSS PLATFORMS AND ITERATIONS.

| **Genos 10B (A100)** | **Fwd** | **RC** | **Fwd/RC Avg.** | **Fwd OR RC** |
|---|---|---|---|---|
| **Benign Set (n=79)** | | | | |
| False Positives (Type I Error) | 4 (5.1%) | 0 (0%) | 18 (22.8%) | 4 (5.1%) |
| True Negatives (Specificity) | 75 (94.9%) | 79 (100%) | 61 (77.2%) | 75 (94.9%) |
| **Splicing Set (n=49)** | | | | |
| True Positive (Sensitivity) | 22 (44.9%) | 4 (8.2%) | 29 (59.2%) | 24 (49%) |
| False Negative (Type II Error) | 27 (55.1%) | 45 (91.8%) | 20 (40.8%) | 25 (51.0%) |
| **Accuracy** | **75.8%** | **64.8%** | **70.3%** | **77.3%** |
| **Genos 10B (RTX PRO 6000)** | **Fwd** | **RC** | **Fwd/RC Avg.** | **Fwd OR RC** |
| **Benign Set (n=79)** | | | | |
| False Positives (Type I Error) | 4 (5.1%) | 10 (12.7%) | 11 (13.9%) | 13 (16.5%) |
| True Negatives (Specificity) | 75 (94.9%) | 69 (87.3%) | 68 (86.1%) | 66 (83.5%) |
| **Splicing Set (n=49)** | | | | |
| True Positive (Sensitivity) | 19 (38.8%) | 13 (26.5%) | 23 (46.9%) | 28 (57.1%) |
| False Negative (Type II Error) | 30 (61.2%) | 36 (73.5%) | 26 (53.1%) | 21 (42.9%) |
| **Accuracy** | **73.4%** | **64.1%** | **71.1%** | **73.4%** |
| **Genos 10B v2 (A100)** | **Fwd** | **RC** | **Fwd/RC Avg.** | **Fwd OR RC** |
| **Benign Set (n=79)** | | | | |
| False Positives (Type I Error) | 0 (0%) | 2 (2.5%) | 0 (0%) | 2 (2.5%) |
| True Negatives (Specificity) | 79 (100%) | 77 (97.5%) | 79 (100%) | 77 (97.5%) |
| **Splicing Set (n=49)** | | | | |
| True Positive (Sensitivity) | 2 (4.1%) | 4 (8.2%) | 2 (4.1%) | 6 (12.2%) |
| False Negative (Type II Error) | 47 (95.9%) | 45 (91.8%) | 47 (95.9%) | 43 (87.8%) |
| **Accuracy** | **63.3%** | **63.3%** | **63.3%** | **64.8%** |

NOTE: SUPPLEMENTARY TABLES IV TO VI ARE SUPPLIED SEPARATELY AS INDIVIDUAL .XLSX FILES OWING TO THEIR LENGTH VIA ZENODO (doi.org/10.5281/zenodo.23784751).

### SUPPLEMENTARY TABLE IV
CLASSIFICATION OF *OPA1* SINGLE-NUCLEOTIDE VARIANTS BY MODEL.

### SUPPLEMENTARY TABLE V
CLASSIFICATION OF *OPA1* VARIANTS WITH MULTIPLE ALTERNATIVE ALLELES.

### SUPPLEMENTARY TABLE VI
CLASSIFICATION OF MULTI-GENE TEST SET VARIANTS BY MODEL.

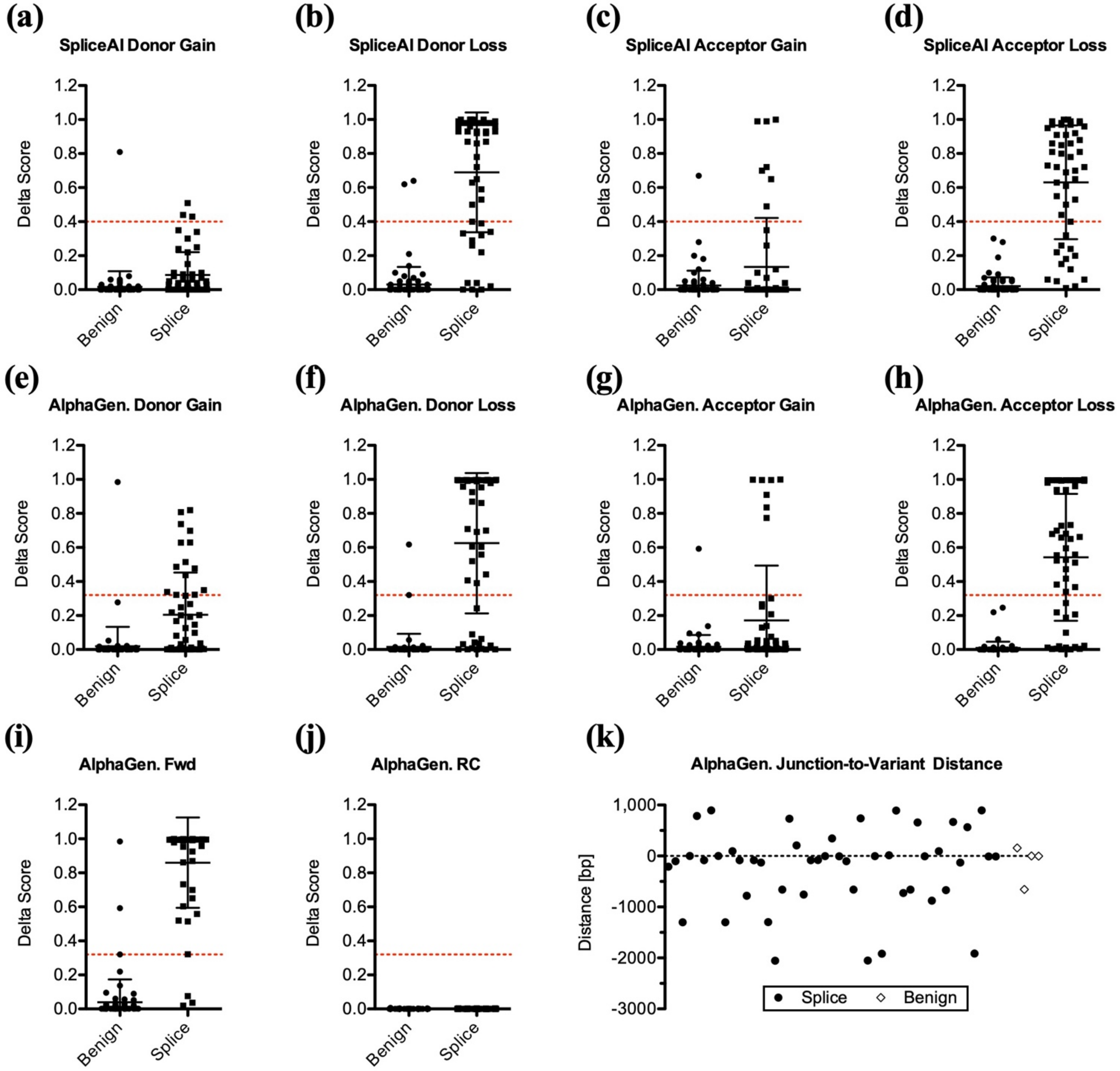


**Supplementary Fig. S1. SpliceAI and AlphaGenome score distributions across individual prediction metrics.** Distribution of delta scores for *OPA1* splicing mutations (Splice) compared to benign sequence variants (Benign). **(a)–(d)** SpliceAI delta scores stratified by acceptor-site gains **(a)**, acceptor-site losses **(b)**, donor-site gains **(c)**, and donor-site losses **(d)**. **(e)–(h)** AlphaGenome splice-site module forward-strand predictions stratified by acceptor-site gains **(e)**, acceptor-site losses **(f)**, donor-site gains **(g)**, and donor-site losses **(h)**. **(i, j)** Only AlphaGenome splice-site module forward (Fwd) strand predictions **(i)** are informative, while reverse-complement (RC) strand predictions remain at baseline **(j)**. Horizontal red dashed lines indicate optimal classification thresholds. **(k)** AlphaGenome splice-junction module junction-to-variant distance predictions in base pairs (bp) relative to the variant position (0), whereby positive and negative values represent downstream and upstream alterations, respectively. Data points represent individual variants (filled circles: *OPA1* splicing set; open diamonds: *OPA1* benign set).

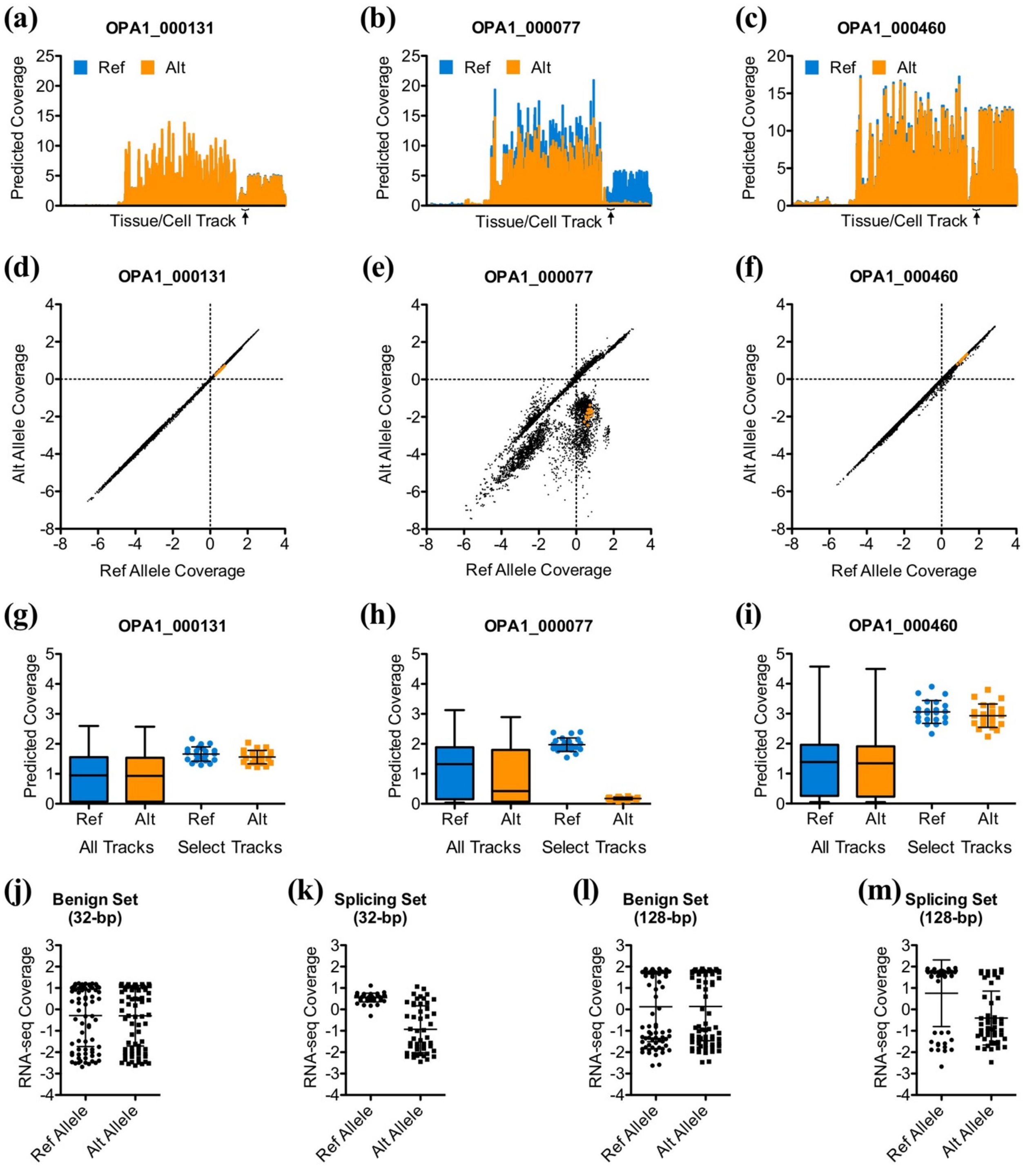


**Supplementary Fig. S2. Borzoi tissue/cell track selection and performance evaluation. (a)–(c)** Predicted RNA-seq coverage across 7,611 human tissue and cell tracks for three representative *OPA1* variants: OPA1_000131 **(a)**, OPA1_000077 **(b)**, and OPA1_000460 **(c)**. Brackets highlight the 22 selected tracks that distinguish reference (Ref, blue) from alternative (Alt, orange) alleles. **(d)–(f)**, Scatter plots showing the correlation of predicted coverage between reference and alternative alleles across all tracks for OPA1_000131 (d, $R^2 = 0.9996$), OPA1_000077 (e, $R^2 = 0.6096$), and OPA1_000460 (f, $R^2 = 0.9981$). **(g)–(i)** Comparison of predicted coverage values between all tracks and selected tracks (orange) for OPA1_000131 **(g)**, OPA1_000077 **(h)**, and OPA1_000460 **(i)**, showing tighter clustering across selected tracks (box plots show median, interquartile range, and 5th/95th percentiles). **(j)–(m)** Predicted RNA-seq coverage of reference and alternative alleles at 32-bp binning for the benign set **(j)** and splicing set **(k)**. A larger bin size (128 bp) increased prediction noise across both the benign set **(l)** and the splicing set, where the effect was most pronounced for the reference allele **(m)**.

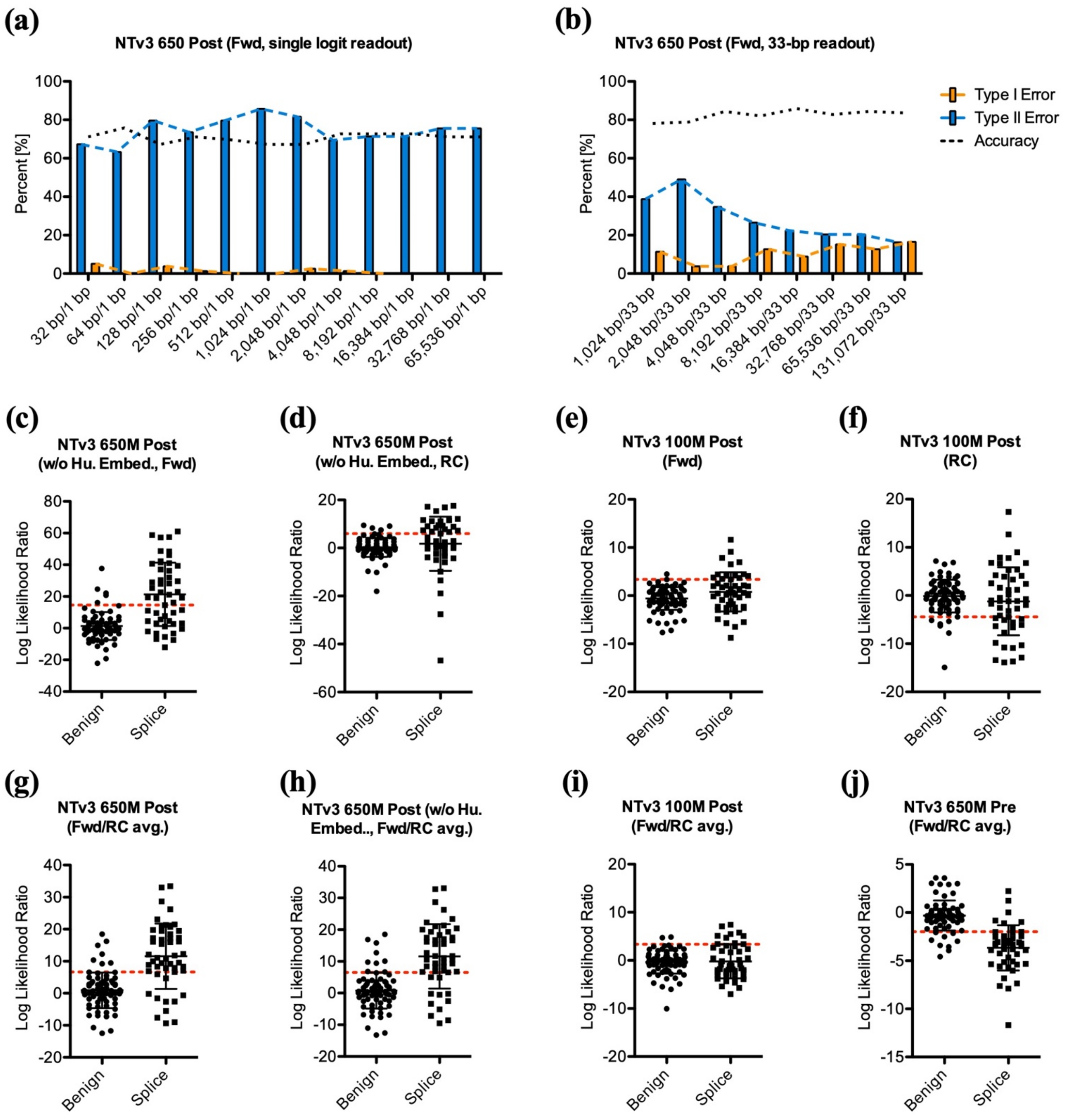


**Supplementary Fig. S3. Context-window scaling, and strand-directionality analysis of Nucleotide Transformer v3 (NTv3) model iterations. (a, b)** Impact of context-window length and readout aggregation on NTv3 650M post-trained model performance metrics (Type I error, Type II error, and overall accuracy). **(a)** Single-logit readout on the forward (Fwd) strand yields high type II error rates (blue bars) regardless of context length (32 to 65,536 bp). **(b)** Aggregating logit scores across a 33-bp window centered on the variant position substantially reduces type II errors and improves overall classification accuracy (black dashed line), peaking at a 16,384-bp context window. **(c)–(f)** Log likelihood ratio distributions for single-strand inferences evaluated on the forward (Fwd) and reverse-complement (RC) strands for NTv3 650M Post without human embeddings **(c, d)** and NTv3 100M Post **(e, f)**. **(g)–(j)** Averaged forward and reverse-complement strand predictions across model variants: NTv3 650M Post **(g)**, NTv3 650M Post without human embeddings **(h)**, NTv3 100M Post **(i)**, and NTv3 650M Pre **(j)**. Horizontal red dashed lines indicate optimal classification thresholds.

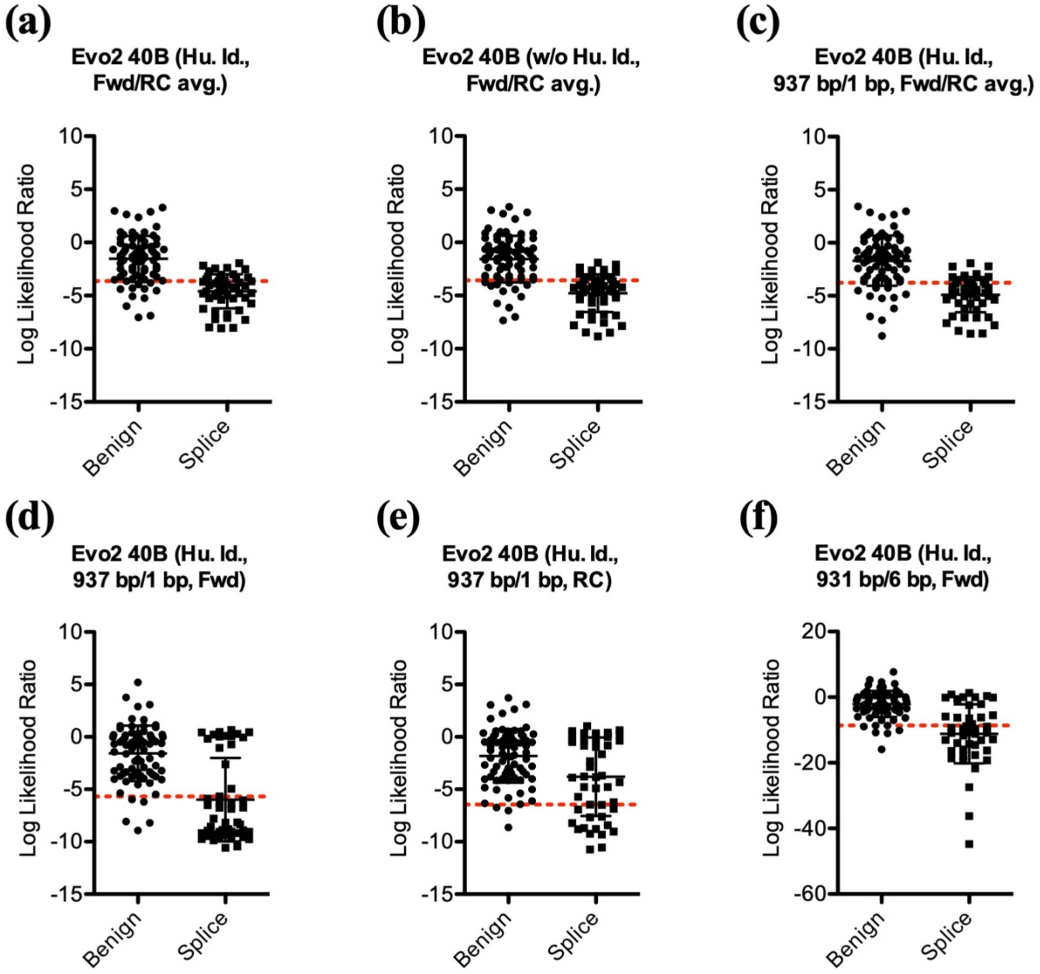


**Supplementary Fig. S4. Evo2 40B performance evaluation and strand-directionality analysis.** Distribution of log likelihood ratios for *OPA1* splicing and benign variants across context lengths, strand directions, taxonomic conditioning, and readout configurations. **(a)** Strand-averaged predictions for the human taxonomic identifier–conditioned model (Hu. Id.), and **(b)** the unconditioned model (w/o Hu. Id.) using a 425-bp context window and 1-bp readout. **(c)–(e)** Impact of extending the context window to 937 bp under human conditioning on strand averaging **(c)** the forward (Fwd) strand **(d)**, reverse-complement (RC) strand **(e)**. **(f)** Logit aggregation across a 6-bp window (931 bp context, 6-bp readout, forward strand), demonstrating signal dilution across variants. Horizontal red dashed lines indicate optimal classification thresholds.

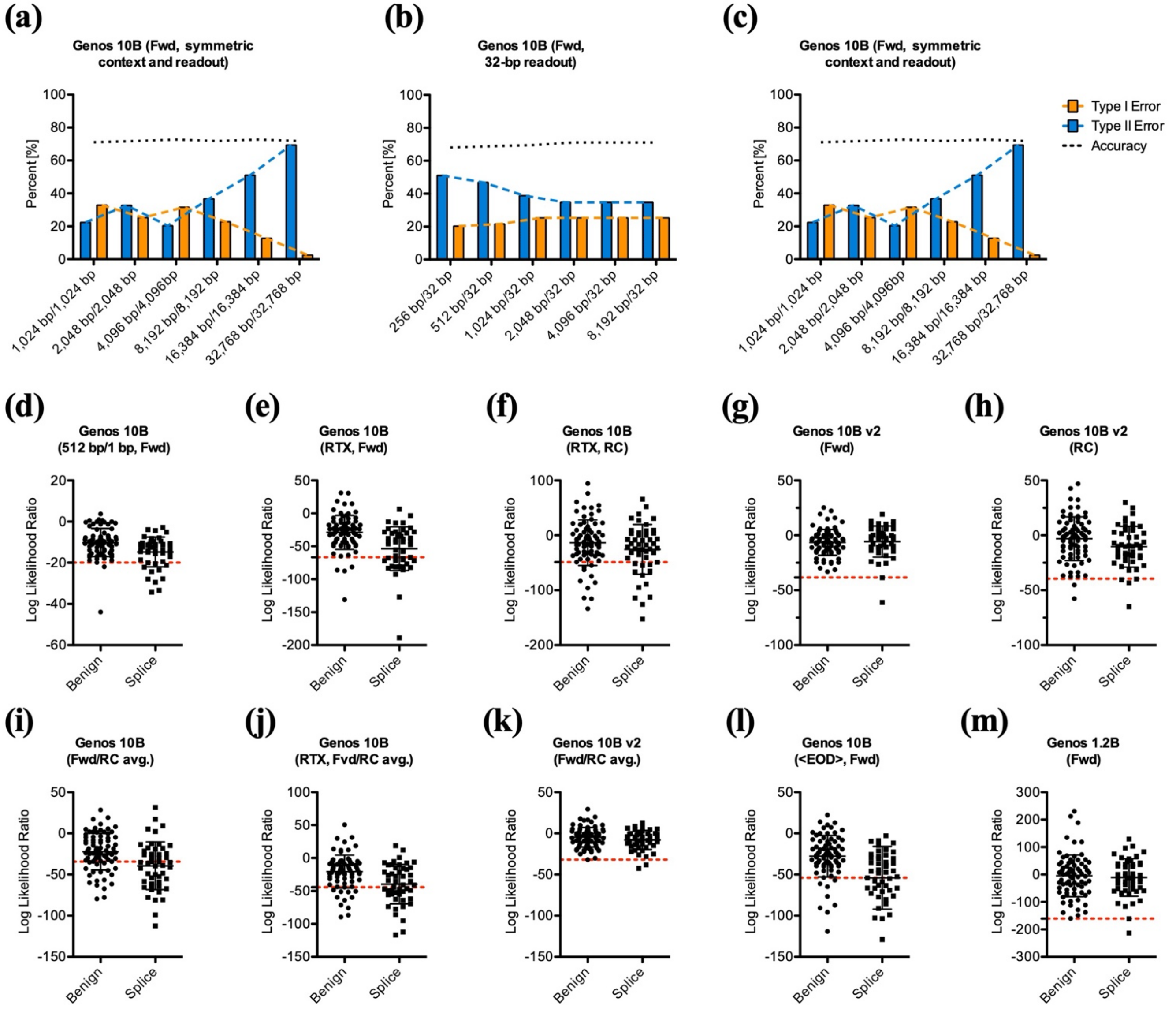


**Supplementary Fig. S5. Genos 10B context-window scaling, model performance evaluation, and platform benchmarking.** **(a)–(c)** Evaluation of performance metrics (Type I error, Type II error, and overall accuracy) on the forward strand across context and readout window configurations, including symmetric scaling **(a, c)** and fixed 32-bp readout scaling **(b)**. **(d)–(m)** Log-likelihood ratio distributions for *OPA1* splicing and benign variants across model iterations, strand directions, hardware platforms, and prompt configurations. **(d)** Baseline Genos 10B single-logit inference (512-bp context, 1-bp readout, forward strand). **(e, f)** Genos 10B predictions evaluated on an NVIDIA RTX Pro 6000 Blackwell (RTX) platform on the forward strand **(e)** and reverse-complement strand **(f). (g, h)** Genos 10B v2 model predictions on the forward strand **(g)** and reverse-complement strand **(h)**. **(i)–(k)** Strand-averaged score distributions for Genos 10B **(i)**, Genos 10B on RTX **(j)**, and Genos 10B v2 **(k)**. **(l)** Incorporating explicit native sequence boundary tokens (<EOD>) into the Genos 10B prompt introduces noise primarily into the benign set. **(m)** Genos 1.2B model predictions. Inferences were executed on an NVIDIA A100 GPU using a 94,208-bp context and 4,096-bp readout window unless specified otherwise. Horizontal red dashed lines indicate optimal classification thresholds. Fwd: forward strand; RC: reverse-complement strand.

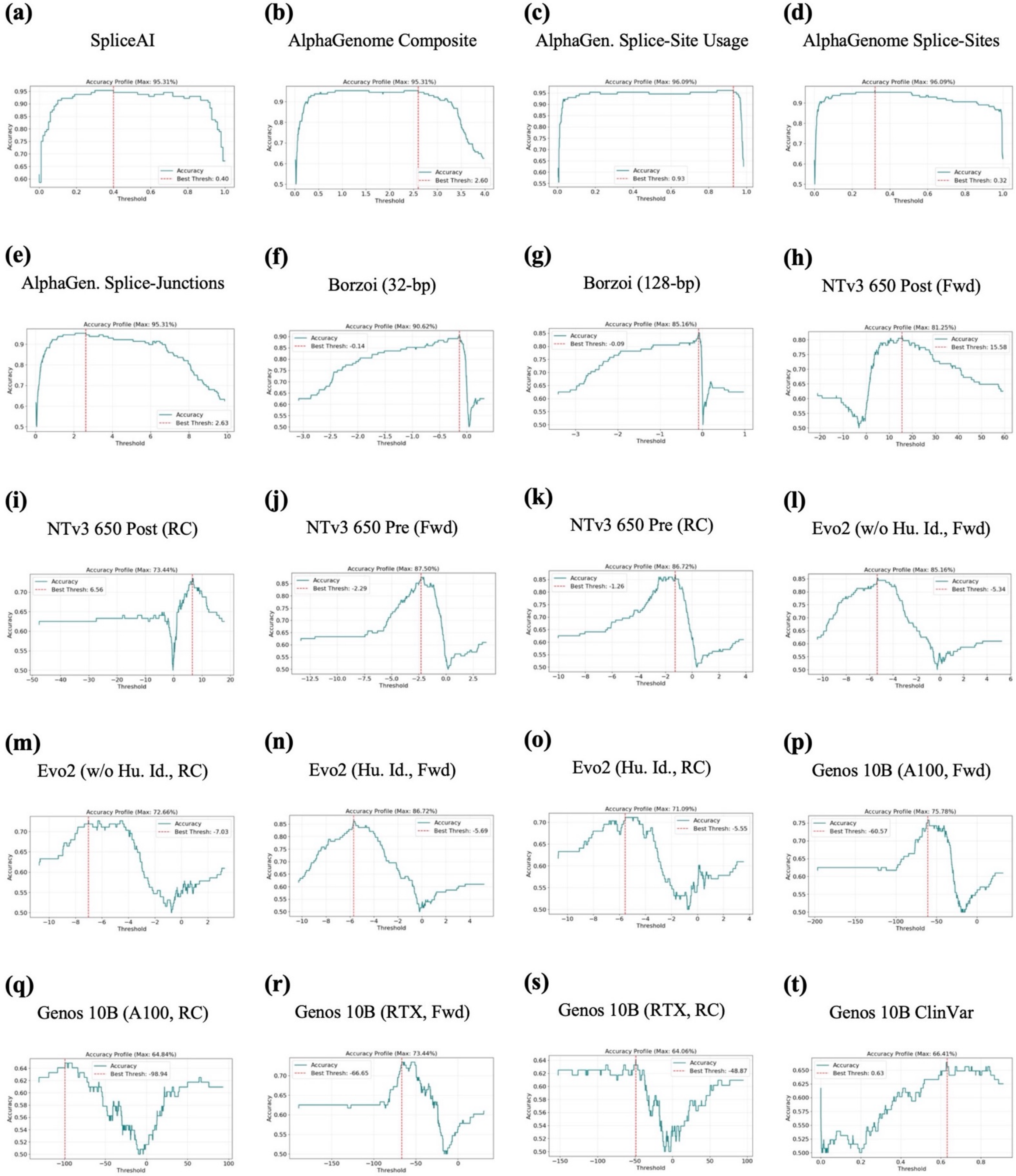


**Supplementary Fig. S6. Determination of classification thresholds optimizing for maximum accuracy while minimizing type I error. (a)–(t)** *OPA1* variant classification thresholds were determined by a one-dimensional grid search across a 10,000-parameter space. Red dashed lines indicate optimal classification thresholds. Fwd: forward strand; RC: reverse complement.

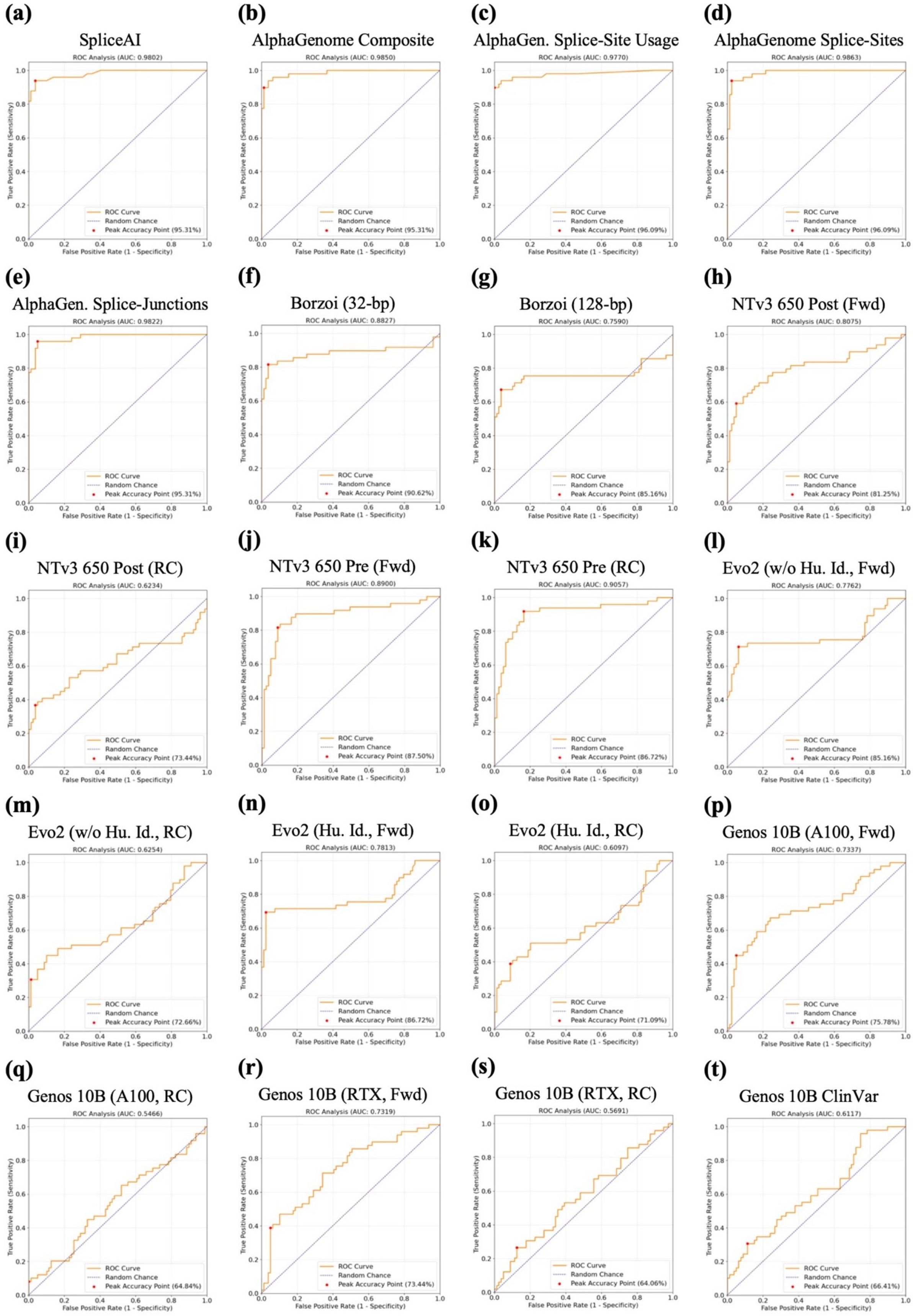


**Supplementary Fig. S7. Area under the receiver operating characteristic (ROC-AUC) curves. (a)–(t)** Individual ROC-AUC curves comparing model performance for *OPA1* splice-variant classification. Orange lines indicate empirical model trajectories; blue dashed lines indicate chance performance (AUC = 0.50). Fwd: forward strand; RC: reverse complement.

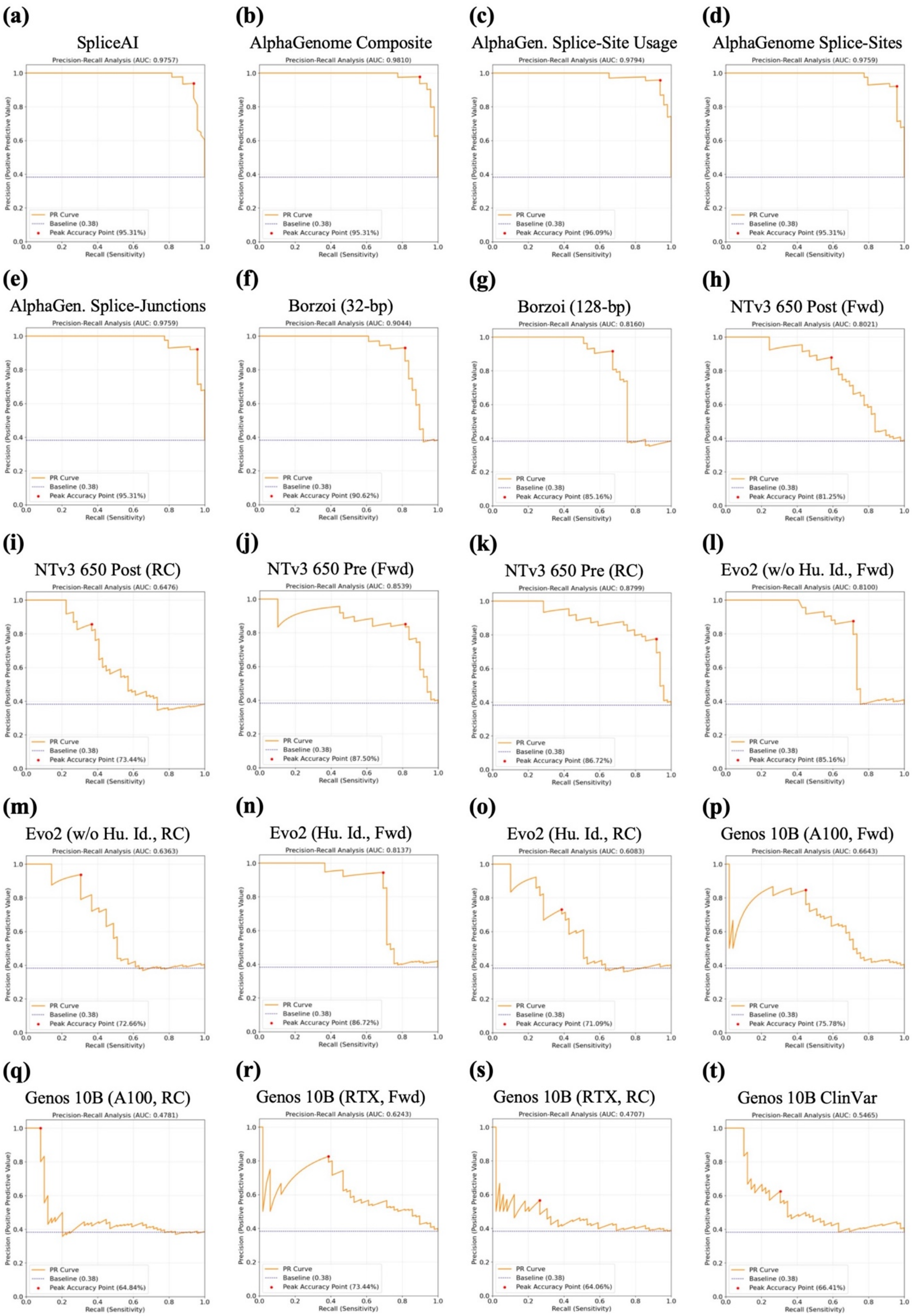


**Supplementary Fig. S8. Precision-recall (PR-AUC) curves. (a)–(t)** Individual PR-AUC curves comparing model performance for *OPA1* splice-variant classification. Orange lines indicate empirical model trajectories; blue horizontal dashed lines indicate baseline chance performance defined by positive class prevalence. Fwd: forward strand; RC: reverse complement.
